# AlphaConformers: Structure-guided sampling enables prediction of multiple protein conformations

**DOI:** 10.64898/2026.08.18.745512

**Authors:** Julie Daniel, Lucas Vitoriano De Queiroz Lira, Diego Javier Zea

## Abstract

Proteins are dynamic molecules capable of adopting multiple conformations. However, AlphaFold2 predominantly generates models around a single conformation, usually representing a ligand-bound state. To address this limitation, we developed AlphaConformers, a structure-guided pipeline that steers AlphaFold2 toward alternative conformations. It is based on the idea that protein structure databases can capture the structural space accessible to members of a protein family. Given a target protein, AlphaConformers retrieves structures from structurally similar proteins. These structures are organized into structure-based alignments and template sets, which are supplied to AlphaFold2 as conformational hypotheses. The resulting models are clustered and filtered, facilitating their analysis. Evaluated on a curated benchmark of 88 proteins with known ligand-bound and unbound conformations, AlphaConformers expanded AlphaFold2 conformational sampling and recovered alternative states missed by AlphaFold2 and other state-of-the-art methods. AlphaConformers ranked first for modelling subtle conformational changes commonly observed between ligand-bound and unbound states. These results show that structural information from protein databases can be leveraged to steer AlphaFold2 toward alternative conformations.

## 1 Introduction

Proteins are the functional building blocks of life, capable of adopting multiple three-dimensional conformations [1, 2]. These conformational changes span a wide range of structural motions, from small side-chain rearrangements and loop remodelling to large domain movements and fold-switching events. In this context, protein conformational changes are often studied by comparing ligand-bound structures, referred to as holo conformations, with ligand-free structures, referred to as apo conformations. In crystallographic apo-holo comparisons, most backbone rearrangements are subtle, with only around 10% of protein families showing RMSD values above 2Å [3]. Consequently, even apparently small conformational differences can be central to protein function.

The capacity to model alternative conformational states is recognised as essential for understanding allosteric regulation, selectivity, and off-target effects in therapeutic design [4–6]. Despite their biological importance, predicting multiple conformational states from sequence alone remains an unsolved challenge [7, 8]. While structure prediction methods such as AlphaFold2 (AF2) [9] have achieved remarkable success [10], they retain a key limitation: predictions tend to converge toward a single preferred conformation, corresponding to the ligand-bound state in 67% of cases [11].

Several strategies have been proposed to address this limitation. A first class of approaches focuses on input manipulation, modifying what is provided to AF2 without altering the model itself [12, 13]. AF-Cluster [14] uses DBSCAN to cluster the input multiple sequence alignment (MSA) and feeds each cluster independently into AF2, based on the hypothesis that distinct sequence clusters encode distinct functional states. Another approach, AFsample3 [15], introduces stochasticity into AlphaFold 3 through random MSA-column masking, thereby perturbing coevolutionary signals and promoting conformational diversity.

A parallel line of work has leveraged structural templates to steer AF2 toward specific conformational states. Heo and Feig (2022) [16] introduced a multi-state modeling protocol that does not use the MSA and instead provides state-annotated templates to AF2. Sala et al. (2023) [17] extended this approach to kinases by automatically retrieving templates filtered by user-defined functional or structural properties. Del Alamo et al. (2022) [12] further demonstrated that combining shallow MSAs with explicit structural templates allows AF2 to predict conformations that would not be sampled otherwise. Together, these studies demonstrate that template selection is a powerful lever for steering AF2 predictions, but previous template-based strategies have largely relied on prior knowledge of relevant conformational states.

Beyond these methods, other approaches adapt AlphaFold-based models for conformational ensemble generation. Rather than introducing variability through the inputs, these methods modify the underlying model architecture to sample conformationally diverse outputs. Among these approaches, AlphaFlow [18] fine-tunes AlphaFold under a flow-matching framework using molecular-dynamics ensembles. In a related direction, BioEmu [19] uses AF2 Evoformer representations within a diffusion-based model trained on molecular simulations and experimental data.

Here, we developed AlphaConformers, a pipeline designed to guide AlphaFold2 toward alternative conformational states. AlphaConformers is based on the idea that the structural space explored by a protein family can contain information about the conformations accessible to individual family members. This idea is supported by the observation that proteins sharing one conformation may also share additional conformations [20]. AlphaConformers therefore leverages structural information already available in databases such as the Protein Data Bank (PDB) [21] and the AlphaFold Database (AFDB) [22] to derive conformational hypotheses that can be supplied to AlphaFold2 and guide its exploration of alternative conformational states.

AlphaConformers retrieves structures from structurally similar proteins available in structural databases. These structures are clustered by structural similarity, and each cluster is then used to construct a structure-based multiple sequence alignment and template set. Both are supplied as inputs to AlphaFold2. To evaluate AlphaConformers, we assembled a curated benchmark of 88 proteins with known apo and holo states. The benchmark was restricted to structures deposited after the AF2 training cutoff, with no structurally similar entries available before that date. We compared AlphaConformers with four state-of-the-art methods spanning sequence-level MSA manipulation and simulation-informed ensemble generation: AF-Cluster, AFsample3, AlphaFlow, and BioEmu. AlphaConformers consistently broadened the conformational space explored by AF2 and was the only method to significantly outperform it for subtle conformational changes. These results show that structural information from protein databases can effectively steer AF2 sampling and improve the recovery of alternative conformational states.

## 2 Results

### 2.1 Overview of the AlphaConformers pipeline

We designed AlphaConformers to expand the conformational space explored by AlphaFold2 (AF2) [9] by taking advantage of the increasing structural information available in protein databases. AlphaConformers can start from a monomeric query structure, either experimental or predicted from sequence. In the first step (Fig. 1A-1), Foldseek [24] is used to retrieve structurally similar proteins from the PDB and the AFDB, generating an initial set of candidate templates and a structure-based MSA. This search is driven by similarity to the query conformation, so known alternative conformations of the retrieved proteins may not always be included. To recover these missing experimentally determined conformations, AlphaConformers uses UniProt identifiers assigned to PDB chains through SIFTS annotations [25, 26] (Fig. 1A-2). These additional representatives of known alternative conformations are added to the initial template set, defining the expanded template set (Fig. 1A-2). The expanded set is then clustered by structural similarity (Fig. 1A-3). For each cluster, AlphaConformers derives a cluster-specific structure-based MSA by retaining only the sequences associated with the structures in that cluster. A corresponding template set of up to four structures is also selected. Both are supplied as inputs to AF2 (Fig. 1A-4; Methods 4.1). To increase sampling within each structural context, AF2 is run multiple times per cluster, yielding 10 predicted structures per cluster (Fig. 1A-5). Because MSA composition and template information can vary across clusters, AF2 confidence scores are not directly comparable across the full ensemble. AlphaConformers therefore includes a scoring step to support model selection and remove low-quality models. Finally, to facilitate the extraction of biological information from the results, the generated models are also clustered (Fig. 1A-6; Methods 4.2).

**Fig. 1:**
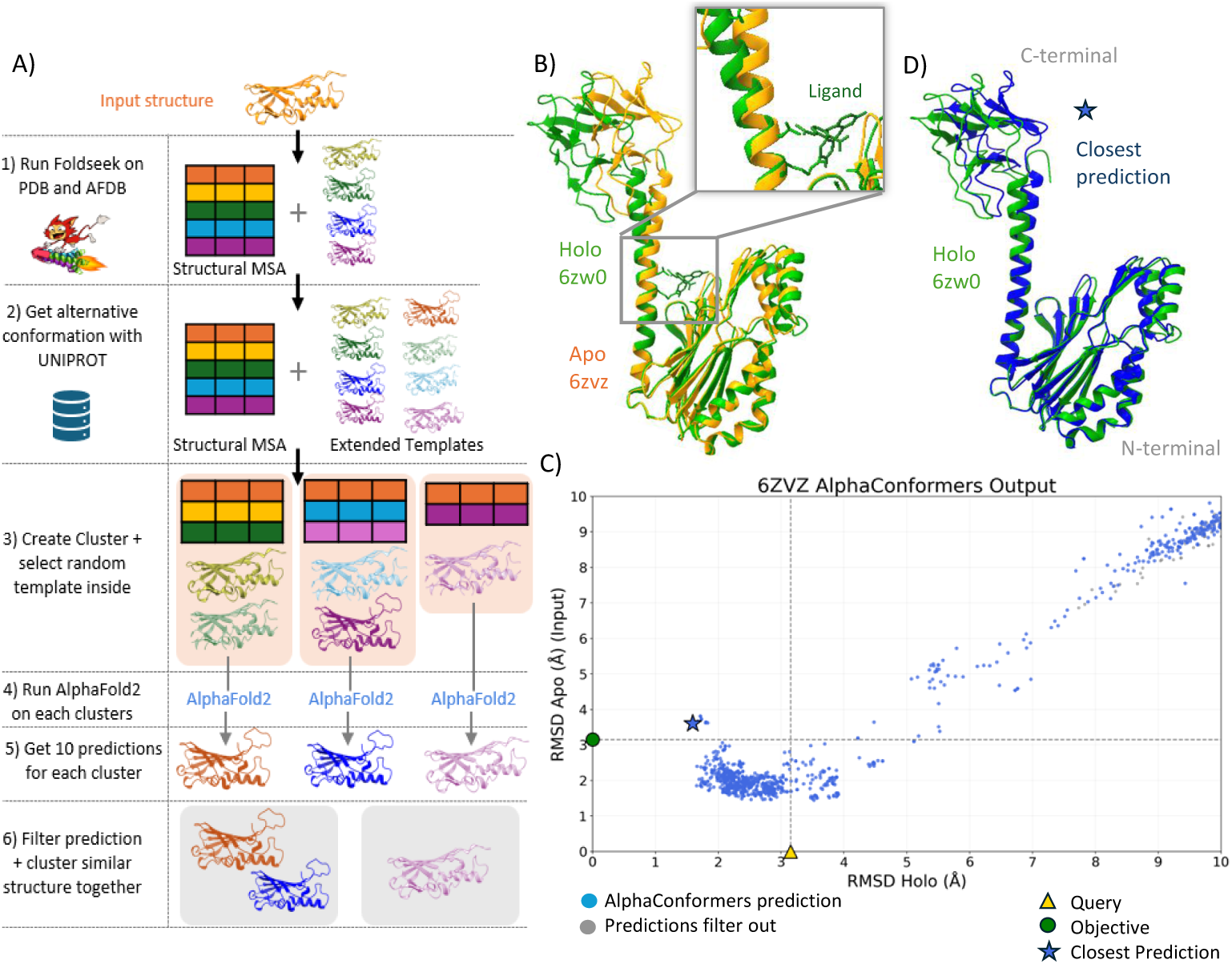
Overview of the AlphaConformers pipeline and illustrative example. **(A)** Schematic of the AlphaConformers pipeline (Methods 4.1). (1) The query structure is submitted to Foldseek for structural similarity search against the PDB and AFDB, yielding a set of structurally similar proteins from which a structure-based MSA is constructed. (2) Alternative conformations of structurally similar proteins are retrieved via UniProt/SIFTS cross-referencing and incorporated into an extended template set. (3) This set is clustered using the Hobohm algorithm (1 Å RMSD cutoff), and up to four representative templates per cluster are provided as input to AF2 alongside the cluster-specific MSA. (4) AF2 is run independently for each cluster, (5) generating 10 predicted models per cluster (2 models x 5 seeds). (6) Predictions are subsequently filtered based on C*β* l-DDT scores from DeepAcNeet[23] and clustered by structural similarity to maximize conformational diversity while reducing the inspection burden (Methods 4.2). **(B)** Illustrative example use for AlphaConformers: protein MJ0548 with structural comparison of the apo (6ZVZ; in orange) and holo (6ZW0; in green) conformations, with the ligand shown in interaction with the holo state, with a apo-holo RMSD of 3.1 Å. **(C)** Visual representation of AlphaConformers output: each point represents a predicted model, positioned according to its RMSD to the apo (6ZVZ) and holo (6ZW0) reference structures, grey point colors correspond to model discarded during the filtering step (6). In this run, the apo structure was used as input (orange triangle), and performance is evaluated with respect to holo recovery (green point). The closest prediction output was represented by a blueu stars. **(D)** Structural superposition of the holo reference structure (in green) and the closest AlphaConformers prediction (in blue), RMSD = 1.7 Å.

**Fig. 2:**
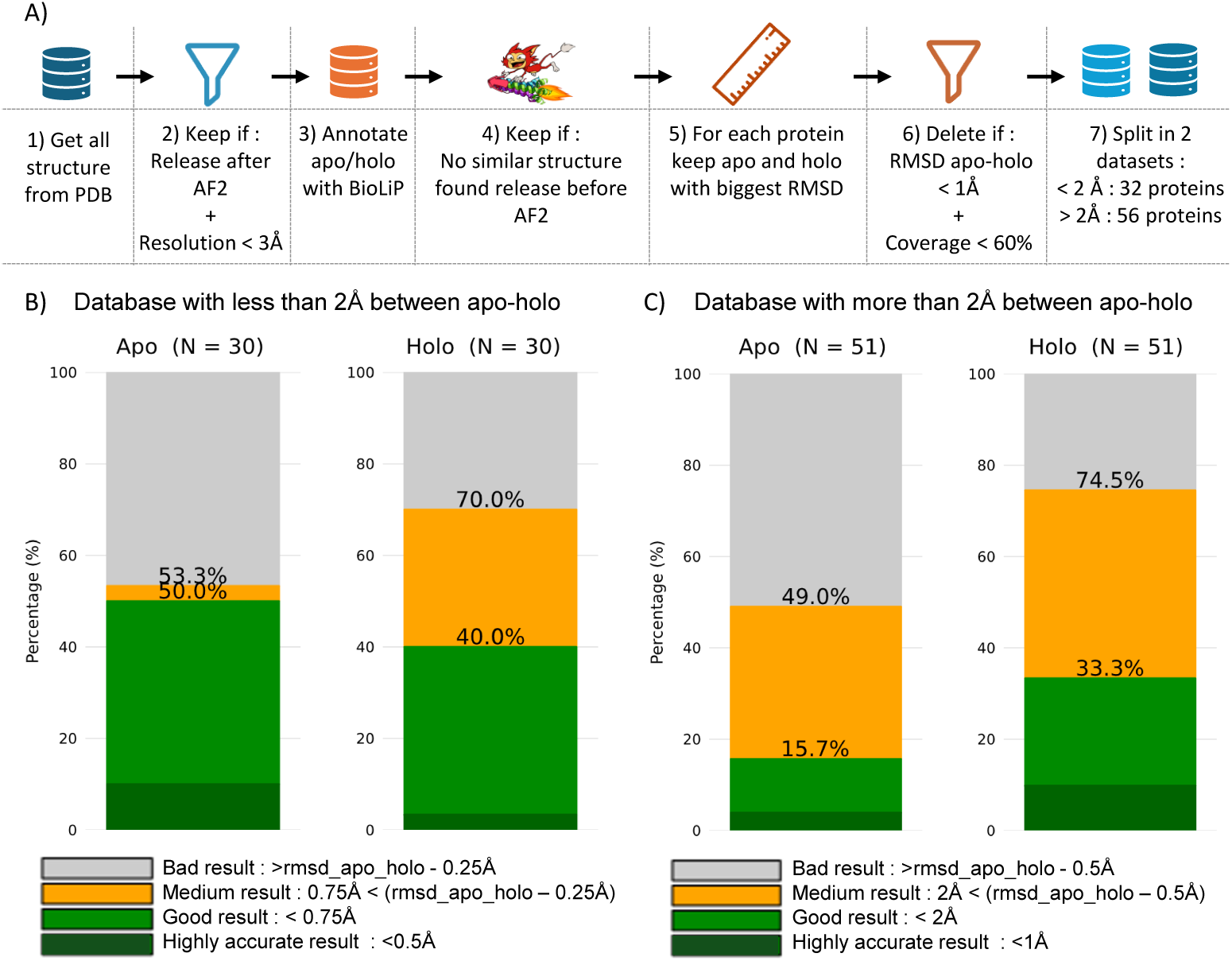
Benchmark dataset construction and AlphaConformers performance in recovering alternative conformational states. **(A)** The benchmark dataset was assembled through seven sequential curation steps (Methods 4.3): (1) all protein structures were retrieved from the PDB; (2) only structures released after the AF2 training cutoff (April 2018) and resolved at 3 Å resolution or better were retained; (3) proteins were annotated using the BioLiP[27] database, structures with a ligand-bound entry in BioLiP were labelled as holo, and those without as apo; only proteins for which both states were available were kept; (4) Foldseek was run on each remaining structure to confirm that no structurally similar entry had been deposited prior to the AF2 training cutoff (details in Methods 4.3); (5) for each protein, the apo-holo pair with the largest RMSD was selected; (6) pairs with an apo-holo RMSD below 1 Å or a sequence coverage below 60% were discarded; (7) the final 88 protein pairs were split into two subsets: a subtle-conformation-change subset (apo-holo RMSD *<* 2 Å, 32 proteins) and a large-conformation-change subset (apo-holo RMSD *≥* 2 Å, 56 proteins). **(B-C)** AlphaConformers was run using either the apo or holo conformation as input and evaluated on its capacity to recover the alternative states. It did not succeed to run on 7 proteins out of the 88, because no similar structure where retrieve by Foldseek. **(B)** Recovery performance on the subtle-conformation-change subset (apo-holo RMSD *<* 2 Å). Predictions are classified as highly accurate (RMSD *<* 0.5 Å), good (RMSD *<* 0.75 Å), or medium (0.75 Å *≤* RMSD *<* apo-holo RMSD *−* 0.25 Å). **(C)** Recovery performance on the large-conformation-change subset (apo-holo RMSD *≥* 2 Å). Predictions are classified as highly accurate (RMSD *<* 1 Å), good (RMSD *<* 2 Å), or medium (2 Å *≤* RMSD *<* apo-holo RMSD *−* 0.5 Å). (Methods 4.4)

**Fig. 3.**
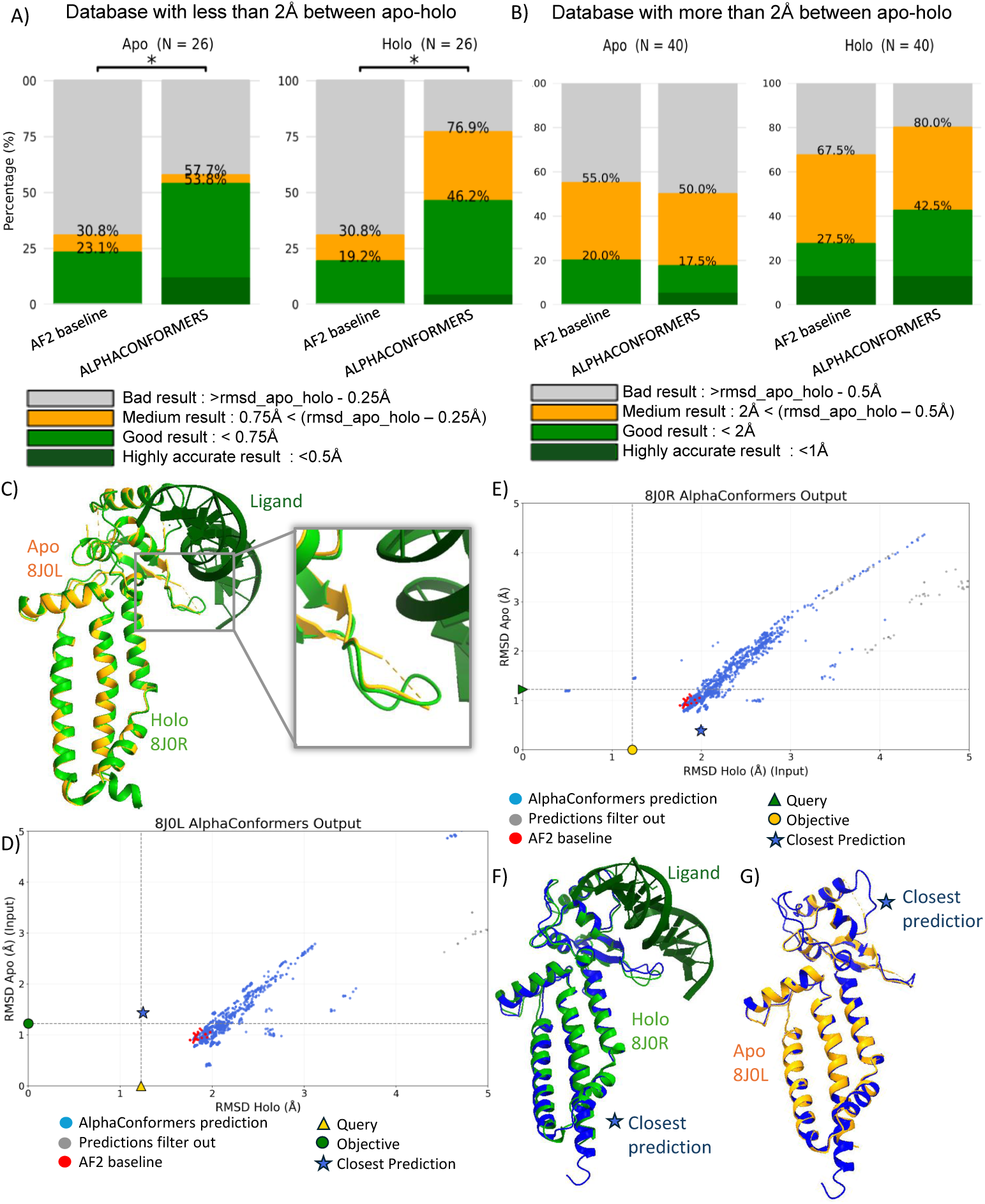
Direct comparison of AlphaConformers and AF2 baseline. A cross-evaluation strategy was applied for AlphaConformers: apo recovery is assessed from runs initiated with the holo structure as input, and holo recovery from runs initiated with the apo structure. AF2 baseline was run with 5 seeds *×* 5 models with dropout enabled. **(A)** Recovery rates on the subtle-conformational-change subset (apo–holo RMSD *<* 2 Å). Statistical significance for both apo and holo recovery was assessed using the exact McNemar test. (Methods 4.5). **(B)** Recovery rates on the large-conformational-change subset (apo–holo RMSD *≥* 2 Å). Statistical significance for holo recovery was assessed using the exact McNemar test **(C)** Illustrative example use for AlphaConformers: the Transcription factor AP-2-alpha with structural comparison of the apo (8J0L; in orange) and holo (8J0R; in green) conformations, with the ligand shown in interaction with the holo state, with a apo-holo RMSD of 1.2 Å. **(D–E)** AlphaConformers output projection for the Transcription factor AP-2-alpha, run with **(D)** the apo conformation (PDB id: 8J0L) as input and **(E)** the holo conformation (PDB id: 8J0R) as input. Each blue point represents a predicted model positioned according to its C*α*-RMSD to both reference structures; grey denote predictions filter out during last step of the the post-processing pipeline (Fig. 1A-6). Red point represents AF2 baseline prediction. Query are represented by triangle, objective as circle and closest prediction with a star **(F)** Structural superposition of the closest AlphaConformers prediction to the holo conformation (blue) with the holo reference structure (green) shown in interaction with the ligand; RMSD = 1.4 Å. **(G)** Structural superposition of the closest AlphaConformers prediction to the apo conformation (blue) with the apo reference structure (orange); RMSD = 0.3 Å

**Fig. 4:**
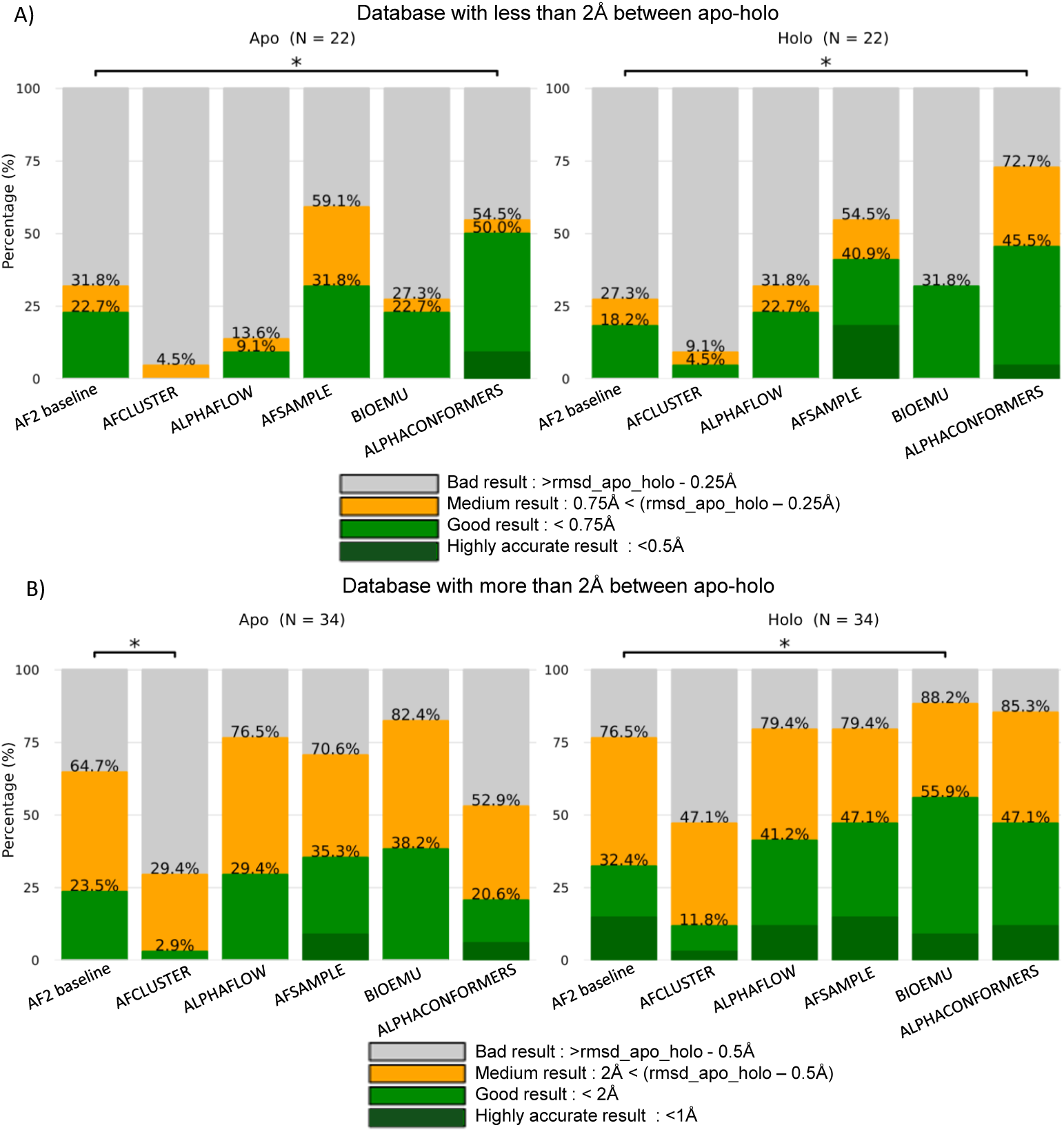
Benchmark comparison of multi-conformational prediction methods on the subtle- and large-conformational-change subsets. Independent recovery rates for the apo and holo conformations are shown across all six methods: AF2 baseline, AF-Cluster, AlphaFlow, AFsample3, BioEmu, and AlphaConformers. For AlphaConformers, a cross-evaluation strategy was applied: apo recovery is assessed from runs initiated with the holo structure as input, and holo recovery from runs initiated with the apo structure. Statistical significance for holo recovery was assessed using the exact McNemar test (Methods 4.5). **(A)** Recovery rates on the subtle-conformational-change subset (apo–holo RMSD *<* 2 Å). **(B)** Recovery rates on the large-conformational-change subset (apo–holo RMSD *≥* 2 Å).

**Fig. 5:**
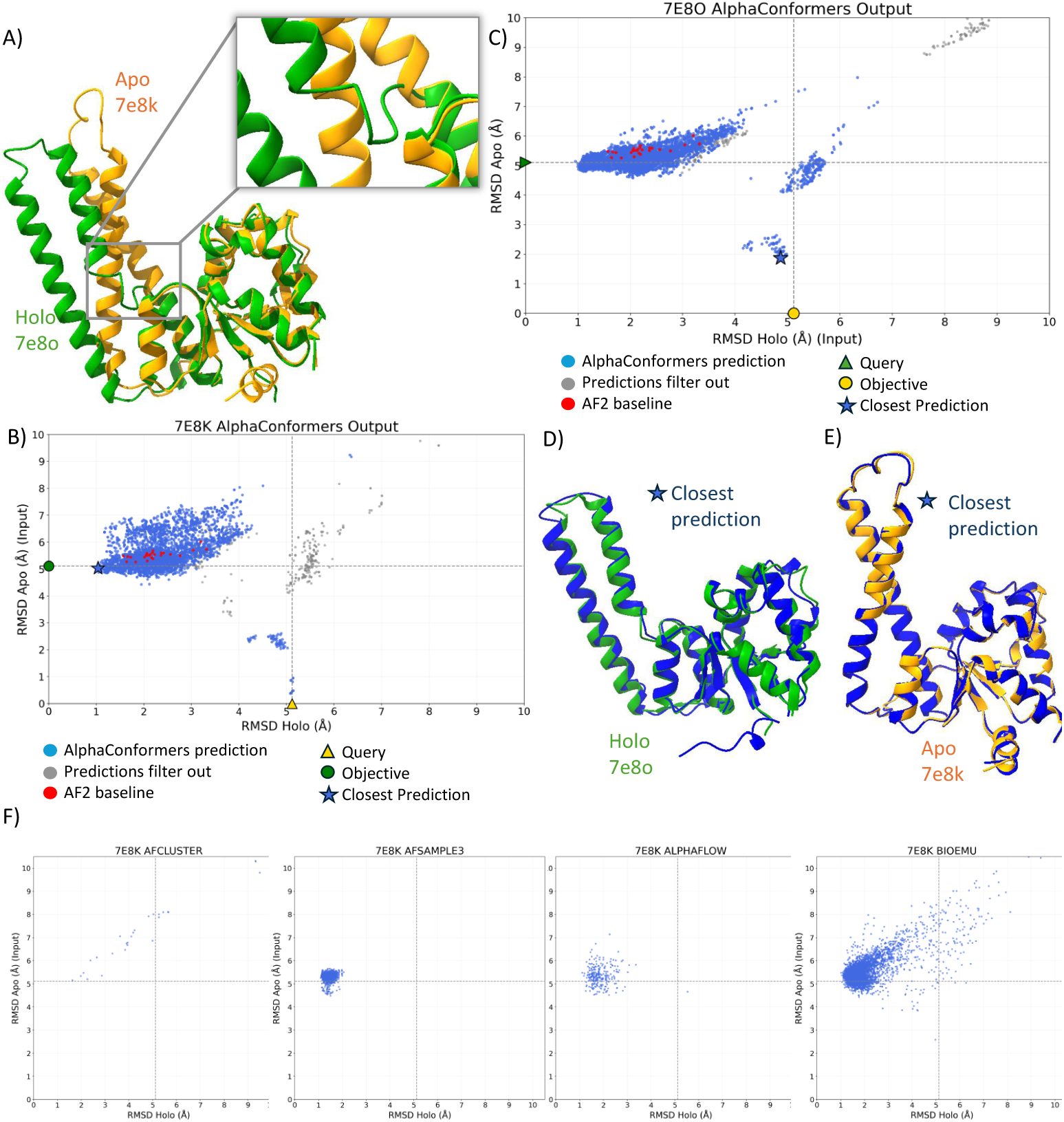
AlphaConformers accurately recovers both conformational states of PbHARP. **(A)** Structural comparison of the apo (7E8K, orange) and holo (7E8O, green) forms, with a RMSD of 5.11 Å and close-up view of the conformational rearrangement at the catalytic site and substrate-binding interface upon pre-tRNA binding. **(B-C)** AlphaCon-formers output projection, run with **(B)** the apo conformation (PDB id: 7E8K) as input and **(C)** the holo conformation (PDB id: 7E8O) as input. Each blue point represents a predicted model positioned according to its C*α*-RMSD to both reference structures; grey denote predictions filter out during last step of the the post-processing pipeline (Fig. 1A-6). Red point represents AF2 baseline prediction. Query are represented by triangle, objective as cercle and closest prediction with a star. **(D)** Structural superposition of the best AlphaConformers prediction (blue) with the experimental holo structure (green), with a RMSD of 1.3 Å. **(E)** Structural superposition of the best AlphaConformers prediction (blue) with the experimental apo structure (orange), with a RMSD of 1.8 Å **(F)** Distribution of predicted models by C*α* RMSD to both the experimental apo and holo structures, shown for AF-Cluster, AlphaFlow, AFsample3 and BioEmu.

We illustrate a specific application of AlphaConformers on protein MJ0548, which both the unbound (apo, PDB id: 6ZVZ) and ligand-bound (holo, PDB id: 6ZW0) conformations experimentally determined, differing by 3.1 Å RMSD (Fig. 1B). Alpha-Conformers was run using the apo conformation as input (represented by a triangle), and performance was assessed by recovering the holo state (represented as a circle). We then computed the RMSD of each predicted structure to both reference conformations, yielding the two-dimensional plot shown in Fig. 1C. The representation shown reflects the output after the full filtering and clustering procedure (Fig. 1A-vi): filtering discarded approximately 2,000 low-quality predictions and eliminated 10 clusters, represented in grey colors. AlphaConformers generates a dense cloud of predictions (in blue) distributed between 1 and 3 Å from both reference conformations, originating near the apo input state and progressively exploring the holo conformational region. The closest prediction (represented by a star) reaches 1.7 Å from the holo ground truth (Fig. 1D). Structural superposition of this best prediction with the holo conformation reveals accurate reconstruction of the N-terminal domain, while the C-terminal domain adopts an intermediate position between the apo and holo states (Fig. 1D). These results demonstrate the ability of AlphaConformers to explore the conformational space from a single input structure and generate predictions in close proximity to the target alternative conformation.

### 2.2 AlphaConformers recovers both apo and holo conformations across a diverse benchmark

To evaluate AlphaConformers rigorously, we constructed a benchmark dataset of 88 protein pairs (apo and holo) through seven sequential curation steps (Fig. 2A). Structures released after the AF2 training cutoff and resolved at sufficient resolution were first retrieved from the PDB (Fig. 2A-1,2). We annotated ligand-bound (holo) and unbound (apo) states using the BioLiP database [27]: structures lacking biologically relevant ligand interactions were labelled apo, and those with at least one BioLiP-annotated ligand interaction were labelled holo (Fig. 2A-3). Each retained structure was then searched against the PDB release before april 2018 to exclude any accession with a structurally similar entry predating the AF2 training cutoff, ensuring that performance estimates reflect genuine generalization rather than memorization [28] (Fig. 2A-4; Methods 4.3). For each UniProt accession, the apo-holo pair exhibiting the largest C*α*-RMSD was selected (Fig. 2A-5), and pairs with an apo-holo RMSD below 1 Å or a sequence coverage below 60% between conformations were discarded (Fig. 2A-6). To assess performance across distinct conformational regimes, the benchmark was partitioned into two subsets based on a 2 Å C*α*-RMSD threshold: a subtle-conformational-change subset of 32 protein pairs (RMSD between 1 and 2 Å) and a large-conformational-change subset of 56 protein pairs (RMSD *≥* 2 Å) (Fig. 2A-7).

AlphaConformers was run independently on both subsets using either the apo or holo conformation as input, and evaluated on its capacity to recover the alternative state. Out of the 88 proteins in the full dataset, AlphaConformers successfully completed on 81; the 7 remaining cases were failed because Foldseek could not identify any structurally similar protein other than the protein query itself, leaving no valid templates for the pipeline. Predictions for the 81 retained proteins were classified according to tiered success criteria adapted to each subset (Methods 4.4).

In the subtle-conformational-change subset, AlphaConformers successfully predicted the apo conformation within 0.75 Å for 50% of proteins, and the holo conformation for 40% (Fig. 2B). In the large-conformational-change subset, the apo conformation was successfully recovered within 2 Å for 15.7% of proteins, while holo recovery reached 33.3% (Fig. 2C). These higher recovery rates indicate that AlphaConformers performs best for subtle apo-holo rearrangements, the regime most commonly observed in crystallographic comparisons [3]. This regime is biologically relevant because even small structural changes can affect protein function [29] and expose cryptic binding pockets relevant to drug discovery [30, 31].

### 2.3 AlphaConformers improves alternative-conformation recovery beyond AlphaFold2 baseline

To assess whether alternative-conformation recovery was attributable to AlphaConformers specifically, we compared the pipeline against an AF2 baseline run with 5 seeds *×* 5 models and dropout enabled, yielding 25 predictions per target (Methods 4.6). This comparison was designed to isolate the contribution of structure-guided sampling from standard AF2 sampling alone. AF2 baseline relies on MSAs extracted from the AFDB server, which were unavailable for 15 of the 81 proteins successfully processed by AlphaConformers. Rather than reconstructing the missing MSAs with a different search procedure, which could introduce an additional source of variation, these 15 proteins were excluded from the analysis. Therefore, all comparisons between AlphaConformers and AF2 baseline reported below are restricted to the 66 proteins for which both methods produced valid outputs.

On the subtle-conformational-change subset, significant improvement is observed for both apo and holo recovery compare to AF2 baseline with 53.8% compared to 23.1% for apo and 46.2% compared to 19.2% for holo (Fig. 3A). On the large-conformational-change subset, AlphaConformers improves holo recovery relative to AF2 baseline by predicted with less than 2 Å for 42.5% of protein compared to 27.5%. For the apo recovery AF2 is a little ahead with 20% compared to 17.5% for AlphaConformers (Fig. 3B).

We illustrate a specific application of AlphaConformers on Transcription factor AP-2-alpha, which both the unbound (apo, PDB id: 8J0L) and ligand-bound (holo, PDB id: 8J0R) conformations experimentally determined, differing by 1.2 Å RMSD (Fig. 3C). AlphaConformers was run twice: once with the apo structure as input (Fig. 3D) and once with the holo structure (Fig. 3E). AF2 baseline predictions cluster at approximately 2 Å from the holo conformation without exploring any other region of conformational space. Whereas AlphaConformers reaches 1.4 Å from the holo reference, when the apo structure is used as input (Fig. 3D).

Structural superposition of the holo reference with the closest AlphaConformers prediction confirms good overall agreement (Fig. 3F). Although this prediction does not cross the apo-holo RMSD threshold and is classified as unsuccessful, it nonetheless places AlphaConformers substantially closer to the holo conformation than AF2 baseline, providing additional structural insight into the ligand-bound state. Conversely, when the holo structure is used as input, AlphaConformers recovers the apo conformation with an RMSD of 0.3 Å, representing a highly accurate prediction (Fig. 3G).

To verify that these gains are not simply attributable to the larger number of predictions generated by AlphaConformers, we ran AF2 mass sampling with an exactly matched number of output models (Supplementary Fig. S1). Recovery rates differed by less than 9% relative to AF2 baseline, with no statistically significant differences observed. Given the substantially higher computational cost of mass sampling and its marginal difference, AF2 baseline was retained as the reference for all subsequent comparisons. Together, these results demonstrate that the conformational gains achieved by AlphaConformers arise from structure-guided sampling rather than from increased prediction volume.

### 2.4 Comparative benchmarking against state-of-the-art multi-conformational prediction methods

To position AlphaConformers within the broader landscape of conformational sampling methods, we performed a systematic benchmark comparison against AF-Cluster [14], AlphaFlow [18], AFsample3 [15], BioEmu [19], and AF2 baseline on our curated dataset. Since several pipelines failed to complete on the subset of the 66 proteins, having MSA form AFDB, due to VRAM, RAM, or GPU time limitations; comparative analyses were restricted to proteins for which all methods produced valid outputs: 22 out of 26 in the subtle-conformational-change subset and 34 out of 40 in the large-conformational-change subset.

All methods were evaluated under identical conditions using the same metrics and model selection criteria (Methods 4.4), with each method run using the optimal parameters reported in the respective publications (Methods 4.6). Statistical significance of pairwise performance differences was assessed using exact McNemar tests [32], as detailed in Methods 4.5.

On the subtle-conformational-change subset, AF-Cluster showed the lowest performance; it did not succeed in recovering the apo conformation for any tested proteins and succeeded for the holo for only 4.5%, behind the AF2 baseline that got 22.7% for apo and 18.2% for holo (Fig. 4-A). AlphaFlow ranked fifth for recovering the apo for 9.1% cases and fourth for the holo with 22.7%. BioEmu, showed low performance on subtle conformational change, recovering the apo for 22.7% of proteins and the holo for 31.8%. AFsample3 ranks in second place by recovering the apo for 31.8% of proteins and the holo for 40.9%. Finally AlphaConformers in the first place, recovering the apo for 50% of proteins and the holo for 45.5%(Fig. 4-A). Beyond outright recovery, AlphaConformers also generates the highest proportion of predictions within the apo-holo RMSD minus 0.25 Å threshold, for 54.5% of proteins toward the apo state and 72.7% toward the holo state, ranking first among all tested methods for the holo state (Fig. 4-A). Statistical comparisons reveal that AlphaConformers is the only method to differed significantly from AF2 baseline for both apo and holo states.

We examined the UpSet plots to visualize complementarity between methods (Supplementary Fig. S2A). In the subtle-conformational-change subset, successful recoveries showed small fragmentation across methods, with many targets either recovered by multiple approaches or missed by all. For apo recovery, AlphaConformers succeeded on four proteins that were missed by all other methods, and missed only one case recovered by BioEmu and AFsample3. For holo recovery, the largest method intersection corresponded to four proteins recovered jointly by AlphaConformers and AFsample3, while AlphaConformers missed only one case recovered by BioEmu and AlphaFlow.

On the large-conformational-change subset, AF-Cluster also achieved the lowest overall performance, recovering the apo conformation for only one single protein (2.9%) and the holo conformation for 4 proteins out of 34 (11.8%), falling behind the AF2 baseline (23.5% for apo and 32.4% for holo) (Fig. 4-B). AlphaFlow and AFsample3 showed similar performance, recovering the apo state for 29.4% and 35.3% proteins, respectively, and the holo state for 41.2% and 47.1%. BioEmu led all methods in retrieving the apo conformation for 38.2% of proteins and the holo for 55.9%. AlphaConformers ranked second for holo recovery with 47.1% successes, equal to AFsample3, but ranked fifth for apo recovery with 20.6% successes (Fig. 4-B). Statistical analysis show that for the apo recovery AF-cluster is significantly worse than AF2 baseline. For the holo recovery BioEmu is significantly better than AF2 baseline. We examined the UpSet plots to visualize complementarity between methods (Supplementary Fig. S2B). On the large-conformational-change subset, successful predictions showed limited overlap between methods, indicating substantial complementarity between approaches. For apo recovery, AlphaConformers uniquely recovered two proteins that were missed by all other methods. For holo recovery, the UpSet plot showed a more fragmented pattern, with successful predictions distributed across a larger number of method intersections, most involving only one or two proteins per intersection.

We further analysed the templates found by AlphaConformers, used as input for AF2, to assess how they contribute to prediction success. In examples where AlphaConformers was the only method to recovered the apo conformation, good predictions were associated with templates structure close to the target, extracted from AFDB (Supplementary Fig. S3).

To assess whether increased sampling alone could reproduce the performance of AlphaConformers, we also evaluated AF2 mass sampling, in which the number of AF2 predictions was matched to the number generated by AlphaConformers for each target. Because of its computational cost, this analysis could be completed only on a restricted subset of proteins.

Nevertheless, including AF2 mass sampling did not alter the overall ranking of methods on either the subtle- or large-conformational-change subset (Supplementary Fig. S4).

Finally, AlphaConformers can also be started from an AF2-predicted structure rather than from an experimentally determined one. In this configuration, the starting structure is generated from the same MSA used by the state-of-the-art methods, and AlphaConformers is evaluated on its ability to recover both apo and holo states within a single run (Supplementary Fig. S5A). Overall, this approach performs below the standard experimental-input configuration, occupying an intermediate position between AF2 baseline and standard AlphaConformers (Supplementary Fig. S5B–C), with no statistically significant differences observed between any method pair (Supplementary Fig. S6). These results suggest that the quality of the starting structure influences how effectively AlphaConformers can steer AF2 predictions; when available, experimentally determined structures therefore remain the preferred starting point.

Taken together, these comparisons show that AlphaConformers has its clearest advantage for subtle apo-holo rearrangements. In this subset, AlphaConformers ranked first for both apo and holo recovery and was the only method to significantly improve over the AF2 baseline in both directions. This result indicates that structural information from the PDB and AFDB can steer AF2 toward alternative conformational states that are not reached by standard sampling alone, achieving recovery below 0.75 Å for a substantial fraction of proteins. For large conformational changes, AlphaConformers showed a less pronounced advantage but remained complementary, recovering alternative states missed by all other tested methods while maintaining a computational cost comparable to the most accessible existing approaches (Methods 4.7).

### 2.5 Case study: AlphaConformers accurately predicts the conformational switch of PbHARP, a protein-only RNase P

To illustrate the capabilities of AlphaConformers in a biologically relevant case, we focus on PbHARP (*Planctomycetes bacterium HARP*), a prokaryotic protein-only RNase P (PRORP) enzyme that catalyzes the 5’-end processing of precursor tRNAs (pre-tRNAs), an essential step in tRNA maturation [33]. HARPs are minimal protein-only enzymes composed of a PINc-domain nuclease. Their substrate recognition and catalytic mechanism remained elusive until the crystal structures of PbHARP in its apo form (PDB id: 7E8K) and in complex with precursor tRNA in holo form (PDB id: 7E8O) were determined[33]. PbHARP functions as a homodimer in its biologically active form; however, all structural analyses and AlphaConformers predictions presented here were performed on the monomeric unit. Comparison of these two monomeric structures revealed a striking large-scale conformational rearrangement, with a C*α* RMSD of 5.11 Å between the apo and holo states (Fig. 5A). This conformational transition remodels the catalytic site and facilitates both pre-tRNA binding at the elbow region and 5’-end cleavage (Fig. 5A), making PbHARP a particularly challenging case for conformational prediction.

AF2 baseline Methods 4.3 predictions (red dots) recover the holo conformation with RMSD *<* 2 Å but did not recover the apo (RMSD *>* 5.5 Å), exceeding the apo-holo RMSD boundary (Fig. 5B-C). To confirm that this disadvantage is independent of sampling depth,

AF2 was additionally run in mass sampling mode using the same total number of predictions as AlphaConformers (N = 4,021) and in naive mode with 1 seed and no dropout (Supplementary Fig. S7A). Mass AF2 predictions converged toward the holo conformation with RMSD *<* 2 Å, while naive AF2 predictions remain above 2 Å for the holo state, but both fail to explore the apo conformation, remaining beyond the apo-holo RMSD boundary.

Then AlphaConformers was run independently using either the apo (7E8K) or the holo (7E8O) structure as input, and evaluated on its ability to recover the alternative state. AlphaConformers successfully recovered both the holo conformation (RMSD *<* 2 Å, apo as input; Fig. 5B) and the apo conformation (RMSD *<* 2 Å, holo as input; Fig. 5C). Structural inspection of the best AlphaConformers predictions confirmed that the elbow region was accurately recovered when either the apo or holo structure was used as input, recapitulating the mechanistic insights reported by the original crystallographic analysis [33] (Fig. 5D-E). PbHARP belongs to the large-conformation-change subset, where AlphaConformers generally shows limited apo recovery (22.6%).

Template analysis for PbHARP indicates that successful recovery was not simply driven by templates already close to the target state. When modelling the holo state from the apo input, AlphaConformers produced a model below 2 Å from the holo conformation using a template more than 7 Å from that conformation. Conversely, when modelling the apo state from the holo input, it produced a model 1.9 Å from the apo conformation using a template 5 Å from that conformation. Thus, in both directions, the final model was closer to the target state than the template used in the corresponding run (Supplementary Fig. S7B).

We additionally run the state-of-the-art methods (Fig. 5F), but none of them succeeded to predict the apo state with less than 2 Å. PbHARP illustrates the ability of AlphaConformers to recover alternative conformational states: both states are accurately recovered regardless of which conformation is provided as input, and the pipeline captures the large-scale structural rearrangement associated with pre-tRNA engagement, succeeding where all other state-of-the-art methods tested fail to do so.

## 3 Discussion

AlphaConformers addresses a central limitation of AlphaFold2 (AF2): although AF2 can generate highly accurate structures, its predictions often concentrate within a restricted region of conformational space. We treat alternative-conformation prediction as a structure-guided exploration problem. This approach is based on the idea that the structural diversity observed across structurally similar proteins can provide hypotheses for conformations accessible to a target protein. This rationale is particularly direct when proteins are known to share one conformation, since they may also share additional conformational states [20]. Thus, structural information available across a protein family can be used to define conformational hypotheses for AF2. AlphaConformers implements this idea by retrieving structurally similar proteins and incorporating their experimentally observed alternative conformations as templates to steer AF2 predictions.

Evaluated on a carefully controlled benchmark designed to exclude structurally similar entries available before the AF2 training cutoff, AlphaConformers consistently expanded the conformational space explored by AF2 and shifted predictions toward alternative conformations that AF2 alone failed to approach. Compared with AF-Cluster, AlphaFlow, AFsample3, BioEmu, and AF2 baseline, AlphaConformers showed its clearest advantage for conformational changes below 2 Å RMSD, where it was the only method to significantly improve over the AF2 baseline for both apo and holo recovery. In this subset, AlphaConformers uniquely recovered four apo conformations missed by all other methods, and the largest holo-recovery intersection corresponded to four proteins recovered jointly by AlphaConformers and AFsample3.

In the large-conformational-change subset, BioEmu achieved the strongest overall recovery, particularly for the holo state. Nevertheless, AlphaConformers provided complementary information, recovering two apo conformations missed by all other methods. Interestingly, all methods performed better at recovering the holo conformation than the apo conformation. This is consistent with previous observations that AF2 preferentially predicts holo-like conformations [11], a tendency that may also influence AlphaFold-based methods. Consistent with this bias, holo recovery showed greater overlap across methods, with multiple approaches successfully recovering the same proteins. Nevertheless, the overlap sizes remained small, indicating that the methods remained complementary.

AlphaConformers differs from existing conformational sampling strategies by using structural information directly at both the MSA-construction and template-selection stages. This distinguishes it from AF-Cluster and AFsample3, which primarily perturb or partition sequence-level MSA information, and from AlphaFlow and BioEmu, which use simulation-informed ensemble generation. In this sense, AlphaConformers generalizes template-guided AF2 modeling beyond manually selected or state-annotated templates, providing an automated way to leverage the structural diversity available in the PDB and AlphaFold DB for conformational prediction. It therefore offers a distinct and complementary alternative to other state-of-the-art predictors.

AlphaConformers is not without limitations. The method is necessarily constrained by the coverage and quality of structural databses; particularly PDB and AFDB. Performance can also be affected by incomplete retrieval of relevant structures due to Foldseek sensitivity, or by the introduction of noisy templates when search thresholds are relaxed. These limitations may partly diminish as structural databases continue to expand and structure-search methods become more sensitive and selective. Our template analysis indicates that prediction accuracy is not explained by template proximity alone: accurate predictions can arise from templates close to the target conformation, but also from templates located more than 4 Å away. This suggests that even templates distant from the target can provide useful structural information to AF2, although how this information is integrated during prediction remains unclear. This suggests that template selection, structure-derived MSAs, and AF2 inference may jointly contribute to successful recovery, raising the question of how AF2 integrates template information when sampling alternative conformations.

Looking forward, two developments would be particularly valuable. First, extending AlphaConformers to protein-protein interaction systems would broaden its application to conformational changes associated with binding, assembly, and interface remodeling. Second, improved reference-free scoring and ranking methods for structural ensembles would represent a major advance. AlphaConformers generates predictions from distinct structure-derived MSAs and template sets, whose sizes and qualities can vary substantially; as a result, standard AF2 confidence metrics are not always directly comparable across outputs and may not reliably identify the most relevant alternative conformations. More robust structural-model scoring would therefore help distinguish biologically meaningful conformational states from low-quality predictions when no experimental ground truth is available.

In summary, AlphaConformers introduces a structure-guided framework for conformational modelling. By automatically organizing available structural data into structure-derived MSAs and template inputs, this strategy steers AF2 toward alternative conformations. AlphaConformers expands the conformational space explored by AF2 and enables recovery of alternative states missed by the AF2 baseline and other state-of-the-art methods. Its strongest gains are observed for apo-holo transitions below 2 Å RMSD, where Alpha-Conformers recovers subtle alternative states that are often missed by other approaches. This is particularly relevant because such subtle rearrangements constitute the majority of crystallographic apo-holo differences associated with ligand binding.

## 4 Methods

### 4.1 AlphaConformers pipeline

AlphaConformers takes as input a monomeric query protein structure in .pdb or .mmCIF format, it can be an experimental structure or a predicted one. Structurally similar monomeric proteins are identified using Foldseek (v10.0.0+; E-value *<* 1*×*10*^−^*^5^) [24] searched against the full PDB [21] and AFDB [22] (last updated October 2025) (technical details in Supplementary

Methods S0.8). For each hit, UniProt/SIFTS [25, 26] cross-referencing is used to retrieve all additional experimentally determined structures deposited under the same UniProt accession, thereby expanding the initial hit set to include all known conformational states of structurally similar proteins [20]. For all tests and results presented in this paper, any structure sharing the same UniProt accession as the query was excluded from the template ensemble and not used in subsequent steps, ensuring that no data leakage occurs and that the pipeline’s performance is genuinely assessed on structurally similar, rather than identical, proteins. This exclusion is implemented as a user-adjustable hyperparameter and can be modified if desired. The resulting ensemble of structures is then clustered using the Hobohm I algorithm at a 1 Å RMSD cutoff to ensure conformational diversity while controlling redundancy [34] (technical details in Supplementary Methods S0.9). All parameter thresholds presented here were selected following systematic testing, as shown in Supplementary Fig. S8.

For each cluster, a structurally informed MSA is constructed from the cluster members, with sequences aligned with Foldseek. Up to four templates are selected per cluster and provided jointly with the corresponding cluster MSA as inputs to AlphaFold2 (AF2) [9]. Four templates correspond to the maximum number of templates AF2 can handle simultaneously. Because of this limitation, a selection strategy was required. We designed an algorithm that partitions the MSA within each cluster into four parts to identify and select structurally diverse representatives. This approach ensures that the chosen templates are as evenly distributed as possible across the sequence space, capturing the diversity encoded in the MSA using only four structures. For clusters containing only five sequences, the first four are selected directly.

Predictions are generated using ColabFold v1.5.5 [35] with AF2 as the backend (num recycle = 3; num models = 2; use dropout = True; technical details in Supplementary Methods S0.10). To optimize runtime and reduce low-quality outputs, a two-stage execution strategy is applied: each cluster is first run with a single random seed, and the full five-seed run is performed only if the resulting pTM score exceeds 0.6; otherwise, the cluster is skip. This threshold was selected based on systematic testing on our development set of 12 proteins from Saldaño et al. [11], where a cutoff of 0.6 effectively eliminated low-confidence predictions without reducing overall performance. Output prediction are a monomeric unit.

### 4.2 Clustering and reference-free filtering

Following execution of AlphaConformers, hundreds to thousands of predicted structures are generated. To enable efficient interpretation of this large ensemble without experimental ground truth, we developed a four-step post-processing procedure designed to minimize the number of structures requiring manual inspection while preserving conformational diversity in the abscence of ground truth.

The first step defines the expected location of the alternative conformation, grounded in the same structural hypothesis underlying AlphaConformers: if a protein shares one conformation with a structurally similar protein, it is likely to share its alternative conformations as well. All structures in the extended template set (Fig. 1A-2) sharing the same UniProt accession are identified, pairwise RMSDs between them are computed, and the maximum value is retained as the upper bound of the expected conformational displacement range. Any AlphaConformers prediction whose RMSD to the query exceeds this threshold is discarded as lying outside the expected conformational region. This step is conditional: if no structure in the extended template set shares the same UniProt accession, it is skipped and all predictions are carried forward.

In the second step, K-means clustering is applied to all remaining predictions to group structurally similar conformations together, producing 30 clusters (Supplementary Fig. S9A-1 and B; technical details in Supplementary Methods S0.11).

The third step reduces the number of candidate structures using DeepAccNet [23] as a model quality metric (technical details in Supplementary Methods S0.12). AlphaConformers runs AF2 with different cluster-specific MSAs and template sets, so the resulting predictions are not directly comparable using standard AF2 confidence metrics such as pLDDT and pTM. Moreover, predictions close to the target conformation can still receive low pLDDT or pTM scores, meaning that filtering directly on these metrics could discard conformationally relevant models (Supplementary Fig. S9C). We therefore used DeepAccNet, which provides a structure-based quality estimate by predicting a per-residue C*β*-lDDT accuracy map independently of the AF2 input MSA or template context. The mean C*β*-lDDT over all residues was used as the global quality estimate. To avoid discarding relevant predictions that may be locally under-scored, filtering is applied at the K-means cluster level rather than to individual structures: a cluster is removed only when its assigned predictions are consistently low-scoring as a whole. Following threshold optimization on the development set (Supplementary Fig. S9D), any K-means cluster in which more than 75% of assigned predictions fall below the bottom 25% of C*β*-lDDT scores for that system is discarded (Supplementary Fig. S9A-2).

As a final step, agglomerative clustering is applied within each surviving K-means cluster using a pairwise C*α*-RMSD distance threshold of 1.0 Å, yielding compact subclusters of highly similar structures (Supplementary Fig. S9A-3; technical details in Supplementary Methods S0.11). During manual inspection, a single representative structure per subcluster is sufficient to survey the full conformational diversity of the ensemble, substantially reducing the number of structures that must be individually examined.

### 4.3 Benchmark dataset construction

To test AlphaConformers we construct a benchmark free of data leakage relative to AF2’s training set, we extracted from the PDB all structures deposited after April 2018, the established AF2 training cutoff[9], remaining those and with an experimental resolution *≤* 3 Å.

Ligand-bound (holo) and unbound (apo) annotations were assigned using BioLiP[27] ligand interaction data (last updated 25 February 2025), Only protein register under the same Uniprot accession having both apo and holo state were keep. To ensure the absence of similar structure within AF2’s training data, each retained structure was searched against the pre-April 2018 PDB using Foldseek. Any PDB accession yielding a hit with TM-score *>* 0.5 and E-value *<* 1 *×* 10*^−^*^5^ was excluded. For each remaining PDB file, the apo-holo pair of the same Uniprot accession, exhibiting the largest C*α*-RMSD was selected, retaining a single representative pair per accession. Protein pairs with an apo-holo C*α* RMSD below 1 Å were excluded from the benchmark, as a displacement of this magnitude is insufficient to assess model quality. Additionally we add a coverage check between apo and holo couple to me more than 60% otherwise the structural comparison will need be efficient and we couldn’t asses pipeline performance correctly, so protein below that threshold were discarded. The final benchmark comprises 88 unique UniProt accessions.

To assess method performance across distinct conformational regimes, this benchmark was further partitioned into two subsets: a subtle-conformational-change subset, comprising apo-holo pairs with a C*α* RMSD *<* 2 Å, intended to evaluate the recovery of subtle conformational adjustments; and a large-conformational-change subset, comprising pairs with a C*α* RMSD *≥* 2 Å, designed to assess the ability to capture large-scale conformational switches between states. Full technical details are provided in the Supplementary Methods S0.13.

To ensure a more stringent analysis, we constructed an additional dataset with an extra sequence homology verification step. For each protein, we retrieved the sequence and queried the NCBI BLAST web server [36] (details are provided in Supplementary Methods S0.14), applying a filter of 30% minimum identity and 80% minimum coverage. For each hit retrieved, we identified the corresponding PDB entry and checked its release date; any protein with a homologous structure deposited before April 2018 was excluded from the dataset. The resulting dataset comprises 44 unique Uniprot accessions, 17 with a C*α* RMSD *<* 2 Å and 27 with a C*α* RMSD *≥* 2 Å. Results are reported in Supplementary Fig. S10 Since AFsample3 is built on AlphaFold3, it may have been exposed to some structures in our benchmark during training. To ensure a fully unbiased comparison, we apply the same initial filters as described above, but with a modified exclusion criterion: rather than excluding all accessions with any structural hit in the pre-April 2018 PDB, we specify the cutoff data to pre-September 2021. The resulting dataset comprises 16 unique Uniprot accessions, 4 with a C*α* RMSD *<* 2 Åand 12 with a C*α* RMSD *≥* 2 Å. Results are reported in Supplementary Fig. S11.

### 4.4 Evaluation metrics

All predicted structures were evaluated against their corresponding experimental apo and holo structures by computing the C*α* root-mean-square deviation (RMSD) following structural alignment, using the MIToS package (v3.10+) implemented in Julia[37] (technical details in Supplementary Methods S0.15).

Predictions were classified according to tiered success criteria adapted to each benchmark subset. For each method, protein, and target conformation, the model with the lowest RMSD to that conformation was selected, and this RMSD value was used to assign the prediction to a success category. For the large-conformational-change subset (apo–holo RMSD *≥* 2 Å), predictions were classified as highly accurate (RMSD *<* 1 Å to the alternative state), good (RMSD *<* 2 Å), medium (RMSD between 2 Å and the apo-holo RMSD minus 0.5 Å), or unsuccessful (RMSD exceeding the apo-holo RMSD minus 0.5 Å). The 0.5 Å margin ensures that predictions representing to the input conformation, which would fall exactly on the apo-holo RMSD boundary, are not erroneously classified as medium. For the subtle-conformational-change subset (apo-holo RMSD *<* 2 Å), where a uniform 2 Å threshold would be uninformative, a scaled criterion was applied: highly accurate (RMSD *<* 0.5 Å), good (RMSD *<* 0.75 Å), or medium (RMSD between 0.75 Å and the apo-holo RMSD minus 0.25 Å), using a correspondingly smaller margin of 0.25 Å.

An important asymmetry in evaluation design must be noted. Competing methods, AF2, AF-Cluster[14], AFsample3[15], AlphaFlow[18], and BioEmu[19], are sequence-based pipelines that generate conformational ensembles from a MSA without requiring a structural input. They are evaluated on their capacity to recover both the apo and holo states simultaneously within a single run. AlphaConformers, by contrast, requires one known conformation as input and is designed to predict the alternative state. Accordingly, it is evaluated on its capacity to recover the alternative state only: when the apo structure is used as input, performance is assessed against the holo state, and vice versa.

AlphaConformers also offers a configuration in which an AF2-predicted structure is used as input in place of an experimental conformation. In this configuration, the AF2-predicted structure is generated from the same MSA as the competing methods and then supplied to AlphaConformers as the query structure. AlphaConformers is then evaluated on its capacity to recover both apo and holo states within a single run, allowing its performance to be assessed when no experimental starting structure is available (Supplementary Fig. S5A).

### 4.5 Statistical analysis

Pairwise comparisons between methods were performed using exact McNemar tests [32] applied to paired observations on the same targets. For each method pair, only discordant targets were considered informative: those on which the first method succeeded while the second failed (*n*_10_), and those on which the second method succeeded while the first failed (*n*_01_). Under the null hypothesis of no difference in performance between the two methods, *n*_10_ follows a Binomial(*n*_10_ + *n*_01_, 0.5) distribution [38], from which a two-sided exact binomial *p*-value was derived [39].

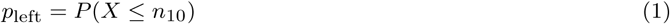

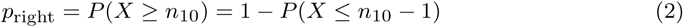

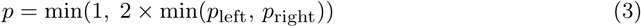

Statistical significance is reported directly on the figures using the following notation: *^∗^ p <* 0.05, *^∗∗^ p <* 0.01, *^∗∗∗^ p <* 0.001; comparisons not reaching statistical significance are left unlabelled.

### 4.6 State of the arts comparisons

To compare AlphaConformers performance we start to compare it to AF2. We run three configuration sharing the same underlying model but differ in the number of predictions generated and the use of dropout. AF2 naive was run using 1 seeds × 5 models (5 predictions total) and without dropout. AF2 baseline was run using 5 seeds × 5 models (25 predictions total) and with dropout. AF2 mass sampling was also run with dropout and generating a number of predictions matched to the total output of the corresponding AlphaConformers run. For a given protein, if AlphaConformers produced predictions across K clusters, AF2 mass sampling was run with (K × 2) seeds × 5 models, ensuring a complete comparison between the two approaches having exactly the same number of output.

Regarding the state-of-the-art methods, AF-Cluster was run using the implementation of Wayment-Steele et al. (2023)[14], with DBSCAN applied to the sequence-based MSA using default parameters. AFsample3 was run with default parameters as described in the paper[15], generating 300 sampled structures per target. BioEmu[19] was run using the publicly available model weights (v1.2) with 4,000 generation steps. AlphaFlow[18] was run using the publicly available fine-tuned weights with 250 starting noise samples. To ensure a fair comparison, all methods were provided with the same MSA, retrieved from the AFDB version 6 using the corresponding UniProt accession; no additional database access was granted. All predictions were evaluated identically across methods using the metrics described in Methods 4.4.

### 4.7 Computational cost

AF2 baseline and AF-Cluster represent the lightest workloads, each requiring approximately 5 GPU-hours per query for 100 predictions (successfully run on 71 and 70 proteins, respectively). AlphaConformers (successfully run on 81 proteins) requires an additional preprocessing overhead for Foldseek searching and structural clustering (approximately 1 CPU-hour per query, reducible to 20 minutes when parallelized across 10 CPU cores), followed by a comparable series of AF2 predictions (typically 100 clusters per query, approximately 5 GPU-hours total). AlphaFlow (successfully run on 64 proteins) is slightly more demanding at approximately 6 GPU-hours per query and requires substantial RAM, preventing execution on 7 proteins on the IDRIS infrastructure (A100/V100, 16 and 32 GB)[40]. BioEmu and AFsample3 are the most computationally demanding methods, each requiring approximately 10 GPU-hours per query on average and successfully run on 66 and 67 proteins, respectively; for some targets, runtimes exceeded 80 consecutive GPU-hours. The differences across methods reflect varying rates of failed runs on specific targets, and therefore all comparative analyses were restricted to proteins for which all methods produced valid outputs (see Methods 4.4).

AF2 mass sampling was evaluated last, requiring approximately 7 GPU-hours for 100 models per query, requiring more GPU-hours by using complete AFDB MSAs, which are deeper than those created by AlphaConformers that are structure-based or the sample ones from AF-Cluster. AF2 mass sampling could only be run on 42 proteins, results for this method are therefore reported separately in Supplementary Fig. S4. Importantly, the inclusion of AF2 mass sampling does not alter the overall ranking of methods on either the subtle- or large-conformational-change subsets. However, pairwise statistical comparisons recalculated on this restricted dataset show that no method significantly outperforms AF2 mass sampling on any evaluated metric, suggesting that while the ranking is preserved, performance differences between methods become narrower when mass sampling is included.

## Code availability

The AlphaConformers pipeline is made fully available through a public GitHub repository diegozea/AlphaConformers.jl. Reproducibility is further supported by two Zenodo archives: one providing the ColabFold/AlphaFold2 configuration used in this study https://doi.org/10.5281/zenodo.20842530, and another containing DeepAccNet, used for the filtering step https://doi.org/10.5281/zenodo.21192188.

## Data availability

The experimentally determined protein structures analysed in this study are publicly available from the Protein Data Bank. The PDB accession codes, chain identifiers, UniProt accessions, apo-holo annotations, curated benchmark data, and numerical source data underlying the figures and statistical analyses are available in Zenodo at https://doi.org/10.5281/zenodo.21133448. The protein structure predictions generated in this study are also available in the same repository.

## Acknowledgements

This work was supported by the French National Research Agency (ANR) through the SPPICES (Scoring and Predicting Protein Interactions and Conformations based on Evolutionary Signals) project (ANR-24-CE45-0866) and performed using HPC resources from GENCI–IDRIS[40] (Grant 2025-AD010317301).

## Author Contributions

Author contributions are defined according to the CRediT taxonomy. Conceptualization: J.D., L.V. (filtering and clustering post-predictions) and D.J.Z. Methodology: J.D. and D.J.Z. Software: J.D. (v2.0 onwards), D.J.Z. (v1.0), Formal analysis: J.D. Investigation: J.D. and D.J.Z. Data curation: J.D. Visualization: J.D. and L.V. Resources: J.D. and D.J.Z. Validation: J.D. and D.J.Z. Funding acquisition: D.J.Z. Project administration: D.J.Z. Supervision: D.J.Z. Writing original draft: J.D. and L.V. Writing review & editing: J.D. and D.J.Z.

