## Supplementary Information for "AlphaConformers: Structure-guided sampling enables prediction of multiple protein conformations"

### Supplementary Methods

#### S0.8 Foldseek structural search: detailed parameters

Foldseek [24] (v10.0.0+0) was run against two databases: the full Protein Data Bank [21] (PDB; download date 17/10/2025) and the AlphaFold Database v6 [22] (AFDB; download date 29/10/2025). Search parameters: alignment type = structural alignment (Foldseek alignment mode enabled), sensitivity = 1, maximum number of sequences = 1,000,000, E-value threshold = $1 \times 10^{-5}$, prefilter mode = disabled, coverage threshold = not specified, sequence identity threshold = none. Searches were conducted on multiple CPUs.

#### S0.9 Hobohm clustering: implementation details

The Hobohm algorithm [34] was applied to the extended template ensemble using a structural RMSD cutoff of 1 Å as the clustering criterion. RMSD on C$\alpha$ was computed using the MIToS package from Julia [37]. Cutoffs of 0.5, 1.0, 1.5, and 2.0 Å were tested; 1.0 Å was selected based on the optimal balance between prediction performance and computational cost (Supplementary Fig. S8). Hierarchical clustering (Ward linkage) [41] was tested as an alternative and yielded comparable performance at substantially higher computational cost; Hobohm was preferred for large-scale deployment.

#### S0.10 ColabFold prediction parameters

Structure predictions were generated using ColabFold v1.5.5 [35] with AF2 weights, using the following parameters: `num_recycles = 3`, `num_seeds = 5` per cluster, and `template_mode = custom` with templates provided explicitly. Computations were performed on V100 or A100 GPUs with 80 GB RAM and 11 CPU cores on IDRIS[40]. The mean computation time per query was approximately 5 GPU-hours for AF2 predictions (across 100 clusters) plus 1 CPU-hour for Foldseek search and structural clustering, yielding a total wall-clock time of approximately 9 GPU-hours per query. A Singularity image reproducing the exact ColabFold environment used in this study is publicly available on Zenodo https://zenodo.org/records/20842530.

#### S0.11 Clustering in reference free filtering

Clustering was performed in two stages. First, an initial partitioning was carried out using K-means clustering [42] (Clustering.jl [43], v0.15.8; `random_state = 42`, `n_clusters = 30`). Each system was represented as an $N \times 3L$ matrix, where $N$ is the number of structures predicted by AlphaConformers and $L$ is the number of residues, with each row containing the superimposed 3D C$\alpha$ coordinates of all residues. K-means clustering was applied directly to this matrix. Second, a refinement stage was applied in which each K-means cluster was further subdivided into structurally tight subclusters: the full $M \times M$ pairwise C$\alpha$-RMSD matrix was computed for each cluster, where $M$ is the number of predictions assigned to it, and used as input to agglomerative hierarchical clustering with average linkage [44] (also via Clustering.jl), using a distance threshold of 1.0 Å.

#### S0.12 DeepAccNet Filtering

DeepAccNet [23] predicts a per-residue C$\beta$ l-DDT accuracy map, the mean over residues was taken as the global model-quality score. Predictions used the NatComm_FA_distance3DBert checkpoint (best.pkl, single model,no 4-model ensemble) with two-body distance features augmented by ProtBert-BFD [45] language-model embeddings (BERT features extracted per structure). Inference was run on an NVIDIA RTX 6000 Blackwell GPU.

#### S0.13 Benchmark dataset, complete filtering pipeline

Proteins entries were downloaded from RCSB PDB in February 2026. Initial filters applied were: deposition date after 30 April 2018 (AF2 training cutoff); experimental method restricted to X-ray crystallography, cryo-EM, or NMR; resolution $\leq 3$ Å; and polymer type limited to protein chains, analysed at the monomer level. UniProt/SIFTS mapping was performed using SIFTS residue-level correspondence files (downloaded November 2025) to map each chain to its UniProt accession. Ligand annotation was carried out using BioLiP [27] (downloaded February 2025): structures with no biologically relevant ligand were labelled apo, and those with at least one BioLiP-annotated ligand interaction were labelled holo. Only UniProt accessions for which at least one apo and one holo structure were available were retained. For each accession, pairwise RMSDs between all available structures were computed, and the apo-holo pair with the largest RMSD was selected. Pairs with an apo–holo RMSD below 1 Å or a sequence coverage below 60% were discarded.

Each retained structure was then searched against a pre-2018 PDB snapshot using Foldseek [24] with the following parameters: structural alignment mode, sensitivity = 1, maximum number of sequences = 1,000,000, E-value threshold $< 1 \times 10^{-5}$, prefilter disabled. Any accession yielding at least one hit with TM-score $> 0.5$ and E-value $< 1 \times 10^{-5}$ was excluded to ensure that no structurally similar entry was available to AF2 during training. The final benchmark comprises 88 unique UniProt accession pairs, split into a subtle-conformational-change subset (apo-holo RMSD $< 2$ Å) and a large-conformational-change subset (apo-holo RMSD $\geq 2$ Å).

An additional held-out test set was constructed using the same curation procedure but with a more stringent deposition cutoff of 30 September 2021, corresponding to the AF3 training cutoff [46], to ensure that AFsample3 predictions are not confounded by memorization of training structures [15]. This subset comprises 15 unique UniProt accession pairs. Results are reported in Supplementary Fig. S11.

#### S0.14 Sequence Homology Filtering

To assess whether the benchmark proteins could have been seen during AlphaFold2 training, we performed a sequence homology search against the PDB for each protein in the dataset. For each entry, the FASTA sequence of the relevant chain was retrieved. The sequence was then submitted to the NCBI BLAST web server using the `blastp` program against the PDB database, requesting up to 500 hits, via the WebBlast.jl package [36]. Hits were retained only if they satisfied both of the following criteria: sequence identity $\geq 30\%$ and query coverage $\geq 80\%$, where identity was computed as the number of identical residues divided by the alignment length, and coverage as the alignment length divided by the query sequence length.

For each retained hit, the corresponding PDB identifier and chain were extracted from the BLAST accession field (format `XXXX_C`). A query protein was excluded from the filtered dataset if any of its hits had been deposited in pre-April 2018 PDB. The entire pipeline was implemented in Julia.

#### S0.15 Structural alignment and RMSD calculation

C$\alpha$ RMSD values between predicted and experimental structures were computed using the Julia programming language (v1.10+)[47] with the MIToS.jl package (v3.10+)[37]. Structural superposition was performed using the superimpose function from the MIToS PDB module, which implements a least-squares optimal superposition of equivalent C$\alpha$ atom pairs using the Kabsch algorithm[48]. Residue-level correspondences between predicted and experimental structures were established retaining only residues present in both structures. The RMSD was then computed over all paired C$\alpha$ atoms following optimal superposition.

### Supplementary Figure

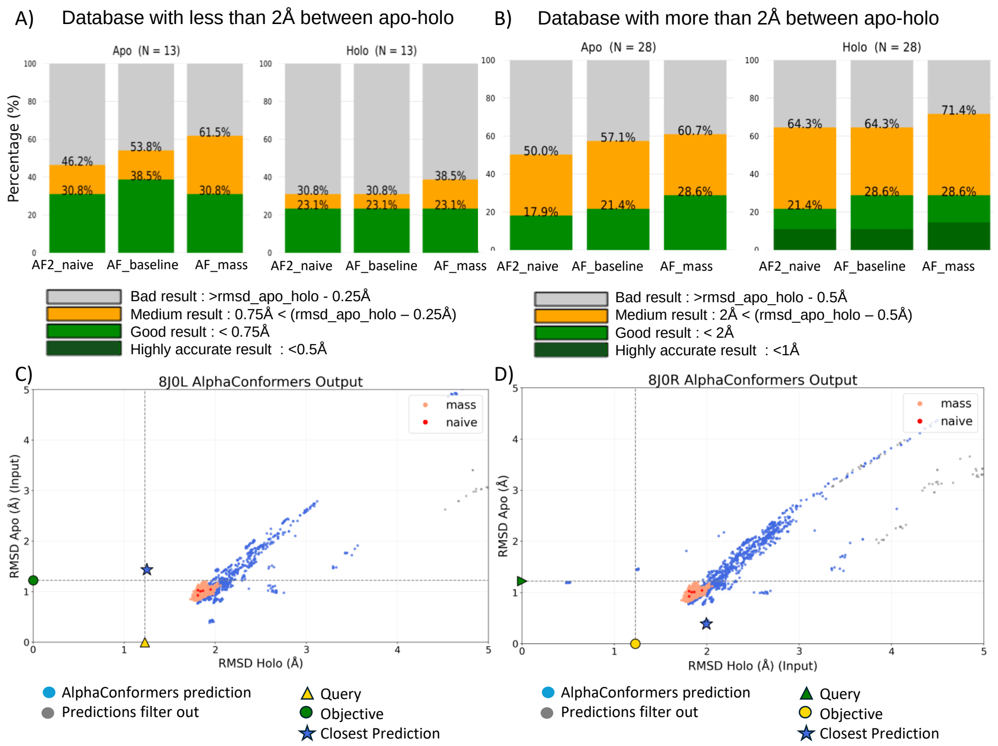


**Fig. S1: Comparison of AF2 sampling strategies** Three AF2 configurations were evaluated: naive (1 seed $\times$ 5 models, no dropout), baseline (5 seeds $\times$ 5 models, dropout enabled), and mass sampling (($K \times 2$) seeds $\times$ 5 models, dropout enabled), where $K$ corresponds to the number of clusters generated by the AlphaConformers pipeline for each target, ensuring an exactly matched number of output models for direct comparison (Methods 4.6). **(A)** Independent recovery rates on the subtle-conformational-change subset (apo–holo RMSD $<$ 2 Å). **(B)** Independent recovery rates on the large-conformational-change subset (apo–holo RMSD $\geq$ 2 Å). Performance is highly consistent across all three AF2 configurations; mass sampling shows a marginal advantage in the medium recovery category, though no differences reach statistical significance. Taken together, panels (A) and (B) demonstrate that increasing the number of AF2 predictions alone does not substantially improve conformational recovery. **(C–D)** Conformational space exploration by AlphaConformers, AF2 mass sampling, and AF2 naive for the Transcription factor AP-2-alpha, run with **(C)** the apo conformation (PDB id: 8J0L) as input and **(D)** the holo conformation (PDB id: 8J0R) as input. Each point represents a predicted model positioned according to its C$\alpha$-RMSD to both reference structures.

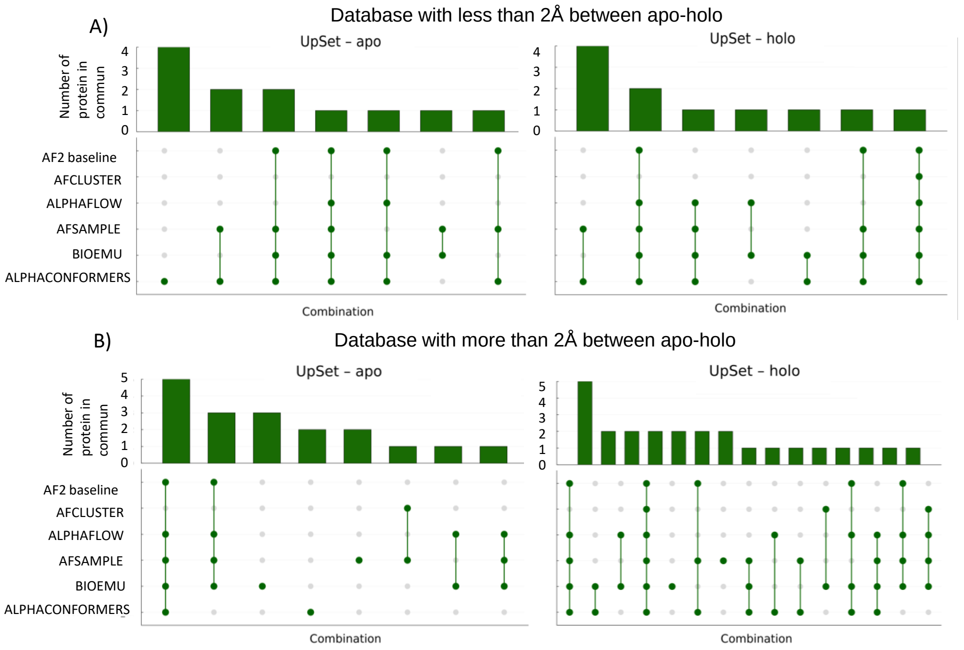


**Fig. S2: UpSet plots of successful conformation recovery across methods.** For each conformation independently (apo and holo), the overlap of successfully predicted proteins across all six methods is shown. **(A)** Subtle-conformational-change subset (apo-holo RMSD $< 2$ Å). For apo recovery, AlphaConformers is the only method to recover 4 proteins not predicted by any other method, while missing only 1 case recovered exclusively by AFsample3 and BioEmu; the remaining 10 proteins are not recovered by any method. For holo recovery, a similar pattern is observed: AlphaConformers misses only 1 case recovered by BioEmu and AlphaFlow, while no method succeeds on the remaining unrecovered targets. **(B)** Large-conformational-change subset (apo-holo RMSD $\geq 2$ Å). In both the apo and holo panels, the largest intersection corresponds to proteins recovered by all methods except AF-Cluster, likely representing targets that are comparatively accessible to most approaches. Beyond this shared subset, successful predictions are distributed across individual methods and small method intersections, indicating substantial complementarity between approaches. This complementarity is particularly pronounced for holo recovery, where successful predictions are more fragmented across methods, with no single approach dominating. Taken together, these results highlight the unique contribution of AlphaConformers to conformational recovery, particularly for small conformational transitions.

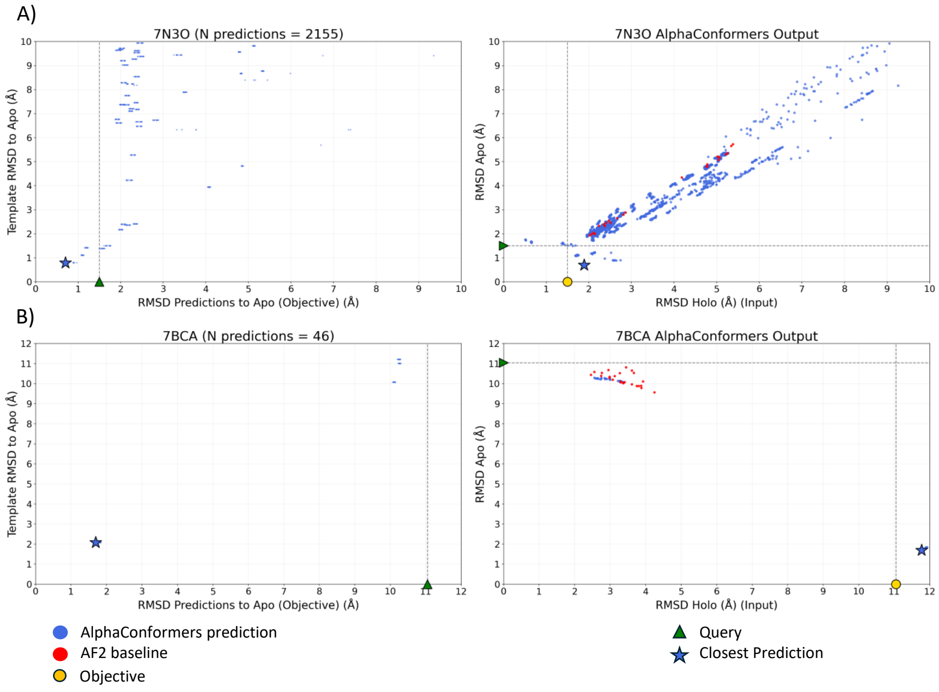


**Fig. S3: Comparison of template proximity and AlphaConformers prediction quality.** Examples in which AlphaConformers is the only method to recover the apo conformation when the holo structure is provided as input. For each example, two panels are shown. Left: RMSD of each predicted model to the apo target conformation plotted against the RMSD of its corresponding template, use as input of AF2, to the apo conformation. Each point represents a predicted model; triangles denote the input (holo) structure and stars denote the closest prediction to the target. Right: two-dimensional projection of the AlphaConformers output, where each blue point represents a predicted model positioned according to its C$\alpha$-RMSD to both reference structures; grey points denote predictions discarded during the post-processing pipeline (Fig. 1A-vi); red points represent AF2 baseline predictions. The input structure is shown as a triangle, the target conformation as a circle, and the closest prediction to the target as a star. **(A)** Cas12k–sgRNA complex, with the holo structure (PDB: 7N3O) used as input, one of the four cases in the subtle-conformation-change subset where AlphaConformers is the only method to recover the apo conformation (PDB: 9GO0) within 0.75 Å. The closest prediction arises from a template within 1 Å of the apo form; an elongated band of predictions is also observed, in which templates at varying distances from the target yield similar prediction RMSDs. **(B)** TrfB transcriptional repressor, with the holo structure (PDB: 7BCA) used as input, one of the two cases in the large-conformational-change subset where AlphaConformers is the only method to recover the apo conformation. A similar pattern is observed, with the best prediction arising from a template at a comparable distance to the target conformation.

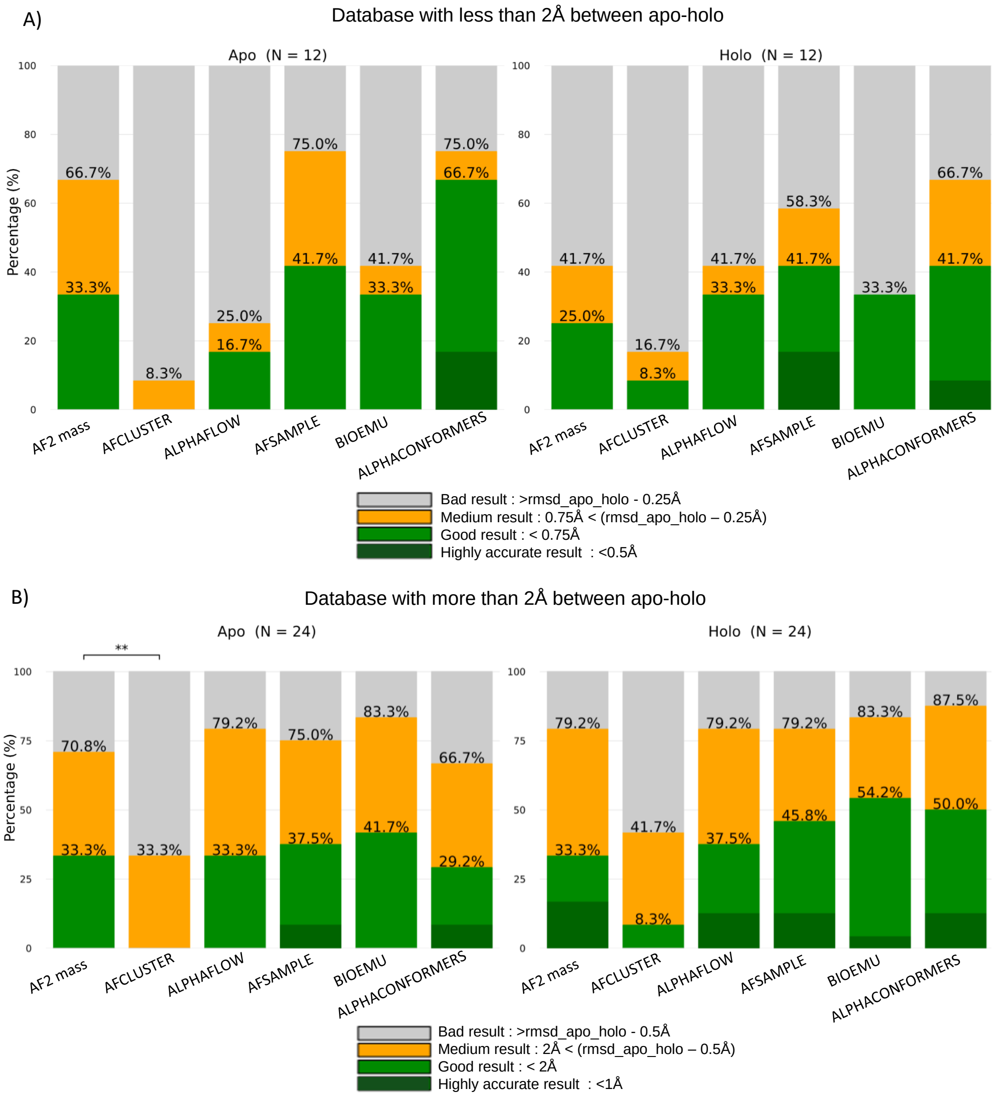


**Fig. S4: Benchmark comparison including AF2 mass sampling. (A)** Independent recovery rates for the apo and holo conformations on the subtle-conformational-change subset (apo-holo RMSD $< 2$ Å). AFsample3 and AlphaConformers still outperform all other methods for both apo and holo recovery. **(B)** Independent recovery rates for the apo and holo conformations on the large-conformational-change subset (apo-holo RMSD $\geq 2$ Å). For apo recovery, methods rank as follows: BioEmu, AFsample3, AlphaFlow and AF2 mass (equal), AlphaConformers, and AF-Cluster. For holo recovery, the ranking is: BioEmu, AlphaConformers, AFsample3, AlphaFlow, AF2 mass, and AF-Cluster. The global ranking doesn't change but only AF-Cluster is statistical worst than AF2 mass sampling

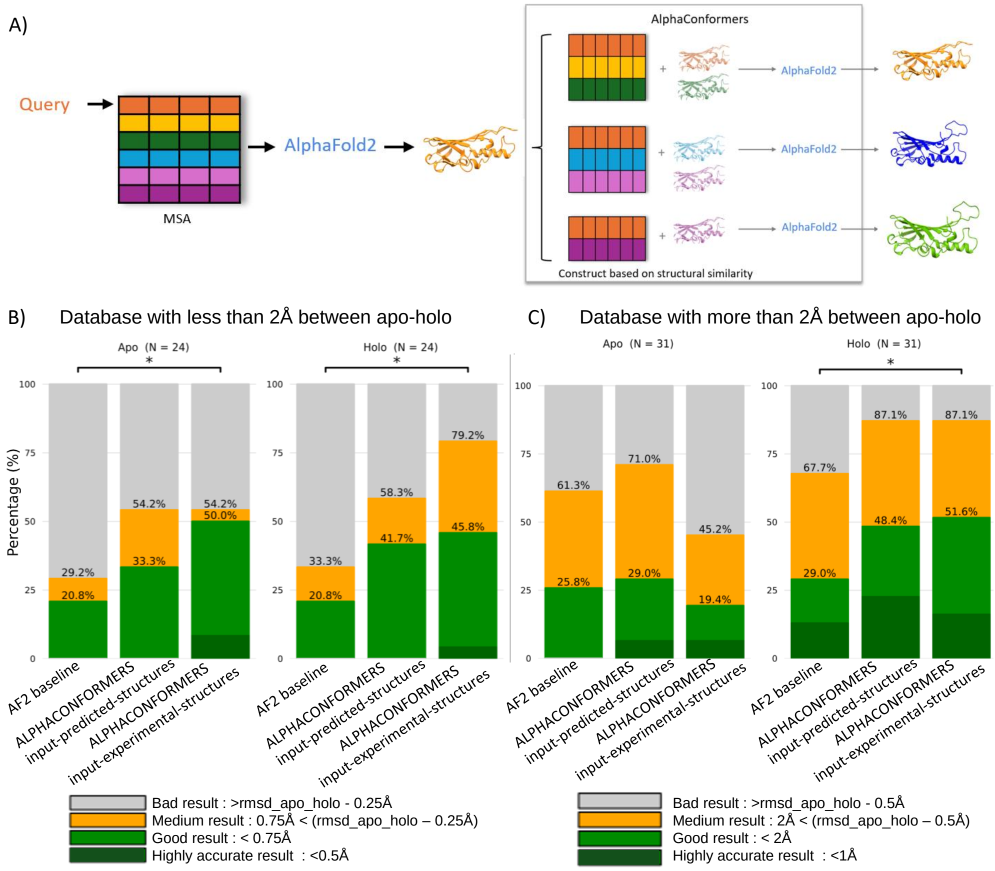


**Fig. S5: AlphaConformers in input-predicted-structures mode. (A)** Extension of pipeline schematic illustrating the input-predicted-structures entry point: a standard MSA is retrive from AFDB server with the UniProt accession, AF2 is run (with 5 models $\times$ 5 seeds), and the highest-confidence prediction (by mean pLDDT) is selected as the input structure for AlphaConformers, which then proceeds identically to its standard operation.**(B)** Recovery rates for the apo and holo states independently on the subtle-conformation-change subset (apo-holo RMSD $<$ 2 Å) compared to AF2 baseline and AlphaConformers in input-experimental-structure mode. Statistical metric calculated with McNemar test (Methods 4.5).**(C)** Recovery rates for the apo and holo states independently on the large-conformation-change subset (apo-holo RMSD $\geq$ 2 Å) compared to AF2 baseline and AlphaConformers in input-experimental-structure mode

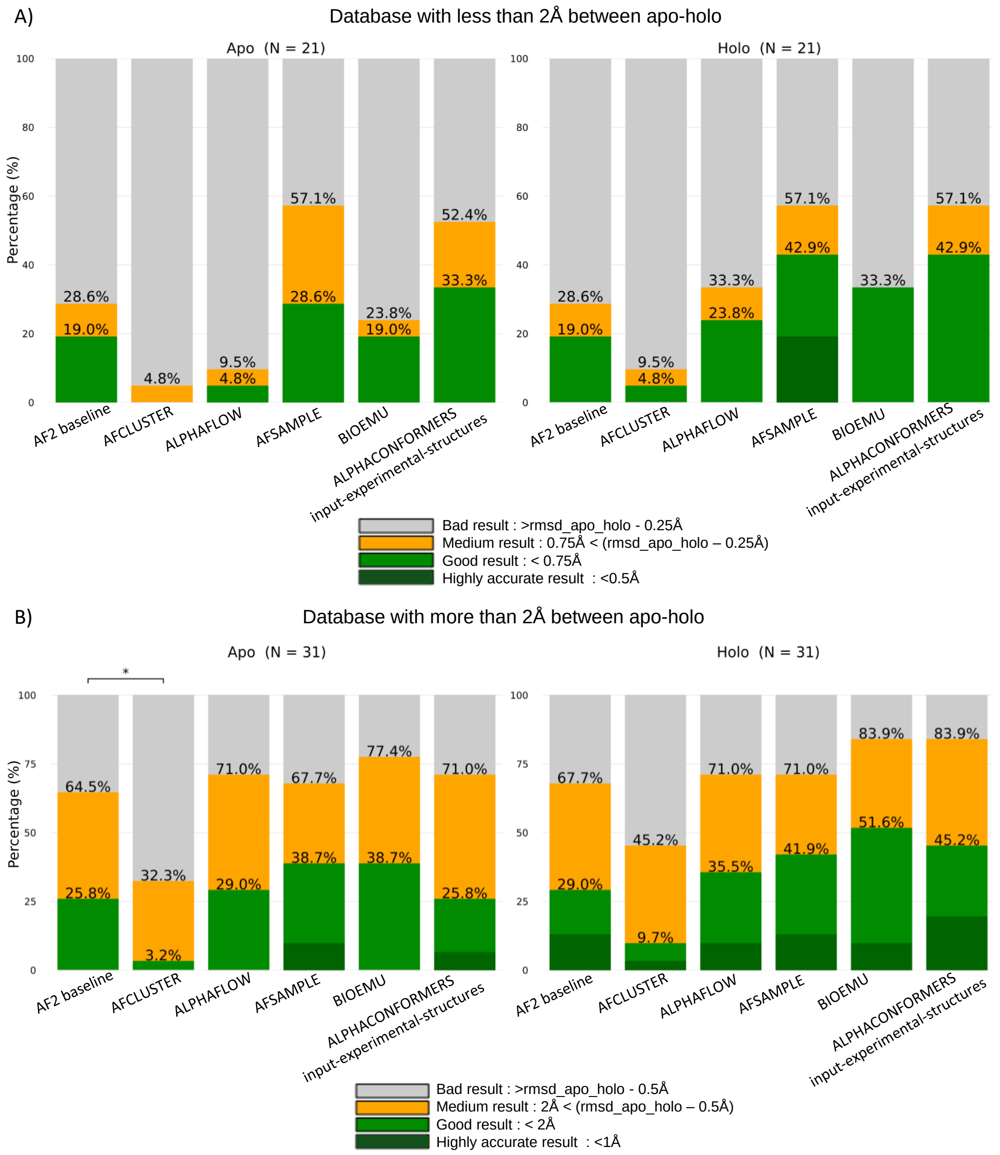


**Fig. S6: Benchmark comparison including AlphaConformers input-predicted-structures mode. (A)** Independent recovery rates for the apo and holo conformations on the subtle-conformational-change subset (apo-holo RMSD $< 2$ Å). AFsample3 and AlphaConformers still outperform all other methods for both apo and holo recovery. **(B)** Independent recovery rates for the apo and holo conformations on the large-conformational-change subset (apo-holo RMSD $\geq 2$ Å). For apo recovery, methods rank as follows: BioEmu, AFsample3 (equal), AlphaFlow, AlphaConformers, AF2 baseline (equal), and AF-Cluster. For holo recovery, the ranking is: BioEmu, AlphaConformers, AFsample3, AlphaFlow, AF2 mass, and AF-Cluster. The global ranking doesn't change but no statistical differences are observed.

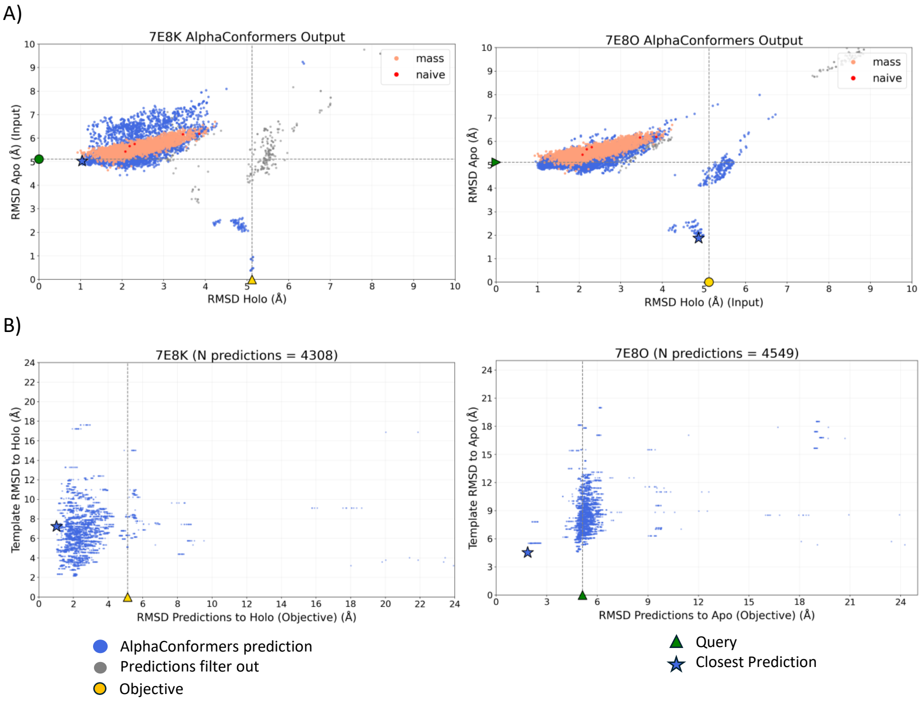


**Fig. S7: Case study on PbHARP: comparison of prediction strategies and relationship between template proximity and prediction accuracy. (A)** Distribution of predicted models according to their C$\alpha$-RMSD to the experimental apo (PDB: 7E8K) and holo (PDB: 7E8O) reference structures, shown for three prediction strategies and two input configurations (apo as input, left; holo as input, right). AlphaConformers predictions are shown in blue; grey points denote models discarded during the post-processing pipeline (Fig. 1A-vi). AF2 naive predictions (5 models $\times$ 1 seed, no dropout) are shown in red, and AF2 mass sampling (($K \times 2$) seeds $\times$ 5 models, dropout enabled) in salmon, where $K$ is the number of clusters generated by the AlphaConformers pipeline ($K = 402$, total predictions $N = 4{,}020$). The blue star marks the closest AlphaConformers prediction to the target conformation; triangles denote the input structure and circles denote the target conformation. **(B)** RMSD of each predicted model to the objective conformation plotted against the RMSD of its corresponding template, use as input of AF2, to the same objective, for both input configurations: apo as input (PDB: 7E8K; RMSD computed to the holo conformation) and holo as input (PDB: 7E8O; RMSD computed to the apo conformation). In both cases, the closest prediction to the target arises from a template located more than 4 Å and up to more than 7 Å from the objective conformation, demonstrating that AF2 extracts and generalises structural information well beyond the immediate vicinity of the provided template.

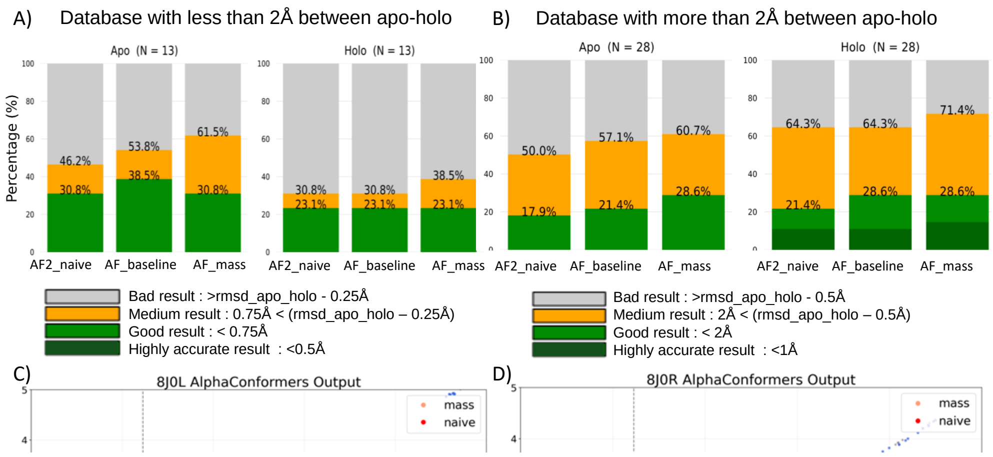


**Fig. S8:** Parameter optimization for database selection, Foldseek[24] search sensitivity, and structural clustering. All tests were performed on a development set of 12 proteins from Saldaño et al.[11], using the apo (unbound) conformation as input. **(A)** Comparison of Foldseek structural similarity searches restricted to the PDB versus searches spanning both the PDB[49] and the AFDB[28]. **(B)** Effect of the E-value threshold on Foldseek search sensitivity, evaluated under three conditions: no threshold (unrestricted), E-value $< 0.1$, and E-value $< 1 \times 10^{-5}$. **(C)** Effect of the Hobohm clustering[34] RMSD cutoff on ensemble diversity, assessed at 0.5, 1.0, 1.5, and 2.0 Å. **(D-E)** Combined parameter analysis isolating the effect of the E-value cutoff with the database fixed to PDB + AFDB: **(D)** results obtained with E-value $< 0.1$; **(E)** results obtained with E-value $< 1 \times 10^{-5}$. **(F)** Summary of all parameter configurations evaluated on the development set. An E-value cutoff of 0.1 yielded bad predictions for 11 out of 12 proteins and so was discarded. Searching across both databases consistently outperformed PDB-only searches. Among clustering cutoffs, 0.5 Å was the least performant, while 1.0 and 1.5 Å yielded similar results. A cutoff of 2.0 Å offered marginal gains only with E-value cutoff filtering NaN, at the cost of increased noise. Based on this analysis, an E-value cutoff of $1 \times 10^{-5}$ combined with an RMSD clustering cutoff of 1.0 Å was selected as the optimal configuration, offering the best balance between computational efficiency and predictive performance. Removing the E-value filter was found to introduce uninformative structures in some cases, substantially increasing computational cost without a performance benefit. Both cutoffs and database use for search are implemented as user-adjustable hyperparameters.

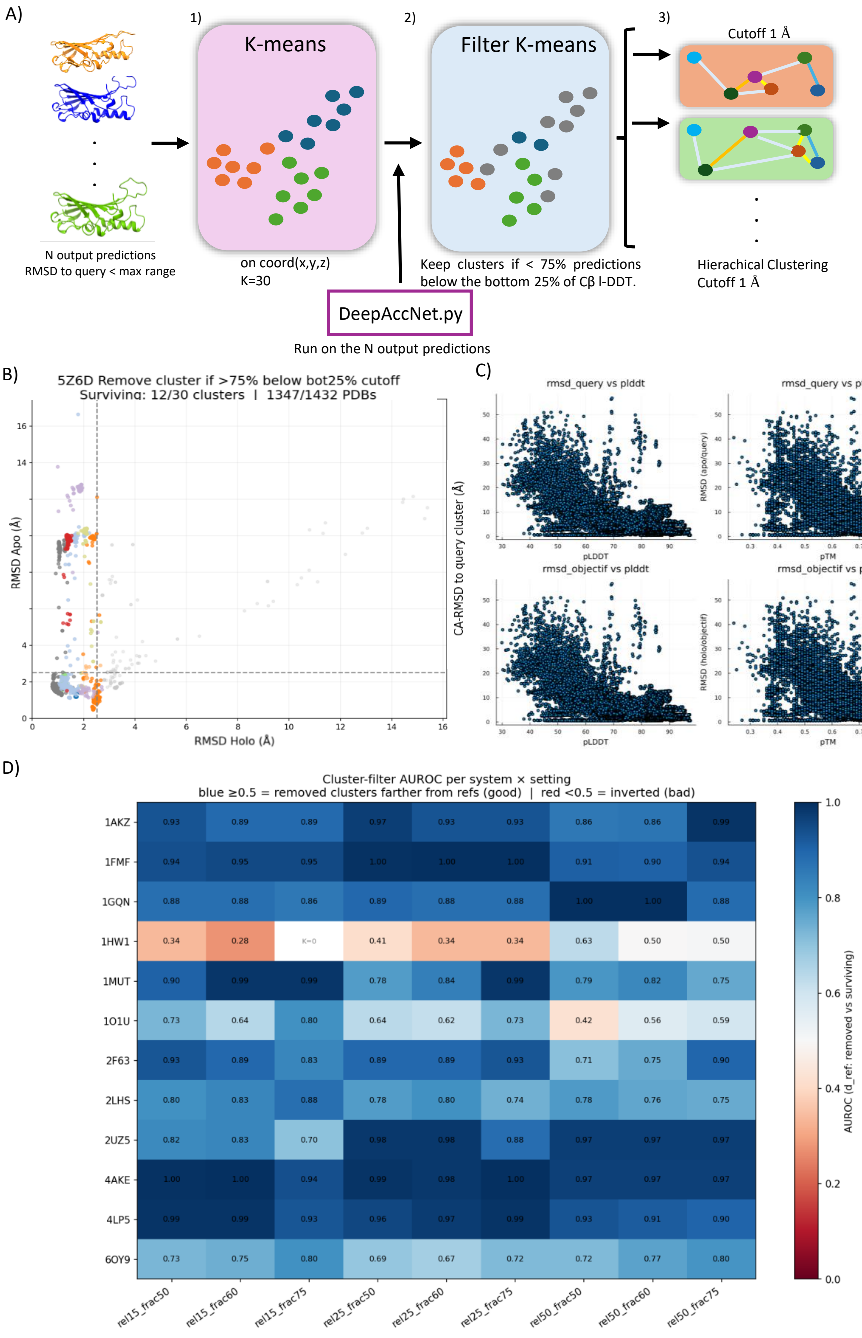
A)
1)
K-means
2)
Filter K-means
3)
Cutoff 1 Å
N output predictions
RMSD to query < max range
on coord(x,y,z)
K=30
Keep clusters if < 75% predictions below the bottom 25% of Cβ l-DDT.
Hierachical Clustering
Cutoff 1 Å
DeepAccNet.py
Run on the N output predictions
B)
5Z6D Remove cluster if >75% below bot25% cutoff
Surviving: 12/30 clusters | 1347/1432 PDBs
RMSD Apo (Å)
RMSD Holo (Å)
C)
rmsd_query vs plddt
rmsd_query vs ptm
rmsd_objectif vs plddt
rmsd_objectif vs ptm
CA-RMSD to query cluster (Å)
RMSD (apo/query)
RMSD (holo/objectif)
pLDDT
pTM
D)
Cluster-filter AUROC per system × setting
blue ≥0.5 = removed clusters farther from refs (good) | red <0.5 = inverted (bad)
1AKZ 0.93 0.89 0.89 0.97 0.93 0.93 0.86 0.86 0.99
1FMF 0.94 0.95 0.95 1.00 1.00 1.00 0.91 0.90 0.94
1GQN 0.88 0.88 0.86 0.89 0.88 0.88 1.00 1.00 0.88
1HW1 0.34 0.28 K=0 0.41 0.34 0.34 0.63 0.50 0.50
1MUT 0.90 0.99 0.99 0.78 0.84 0.99 0.79 0.82 0.75
1O1U 0.73 0.64 0.80 0.64 0.62 0.73 0.42 0.56 0.59
2F63 0.93 0.89 0.83 0.89 0.89 0.93 0.71 0.75 0.90
2LHS 0.80 0.83 0.88 0.78 0.80 0.74 0.78 0.76 0.75
2UZ5 0.82 0.83 0.70 0.98 0.98 0.88 0.97 0.97 0.97
4AKE 1.00 1.00 0.94 0.99 0.98 1.00 0.97 0.97 0.97
4LP5 0.99 0.99 0.93 0.96 0.97 0.99 0.93 0.91 0.90
6OY9 0.73 0.75 0.80 0.69 0.67 0.72 0.72 0.77 0.80
rel15_frac50 rel15_frac60 rel15_frac75 rel25_frac50 rel25_frac60 rel25_frac75 rel50_frac50 rel50_frac60 rel50_frac75
AUROC (d_ref: removed vs surviving)

**Fig. S9: The AlphaConformers reference-free filtering and clustering pipeline. (A)** Schematic overview of the post-processing pipeline. Starting from the $N$ AlphaConformers predictions whose RMSD falls below the expected apo-holo displacement range (Methods 4.2), predictions are processed through three successive steps: (1) initial partitioning by K-means clustering [42] applied to superimposed C$\alpha$ coordinates (Supplementary Methods S0.11); (2) quality-based filtering in which each prediction is scored by DeepAccNet [23], and K-means clusters are discarded when more than 75% of their assigned predictions fall below the bottom 25% of C$\beta$ l-DDT scores for that system (Supplementary Methods S0.12); (3) refinement of surviving clusters by agglomerative hierarchical clustering of the pairwise C$\alpha$-RMSD matrix at a 1 Å distance threshold (Supplementary Methods S0.11). **(B)** Illustrative application to a *Neurospora crassa* protein (apo: PDB id 5Z6D; holo: PDB id 5Z6E); colours denote cluster assignments after filtering. **(C)** Comparison of pLDDT and pTM scores against the C$\alpha$-RMSD to the target and query conformation. Predictions achieving low RMSD values do not systematically receive high confidence scores; conversely, some high-quality predictions in terms of conformational proximity are assigned pTM scores as low as 0.3 or pLDDT scores as low as 40. This decoupling between standard AF2 confidence metrics and conformational accuracy demonstrates that applying a fixed pLDDT or pTM threshold would risk discarding conformationally relevant predictions, justifying the use of a structure-based quality metric independent of the input MSA and templates. **(D)** Selection of the DeepAccNet filtering threshold. Per-system cluster-filter AUROC scores computed on the development set of 12 proteins from Saldaño et al. [11] under varying threshold settings.

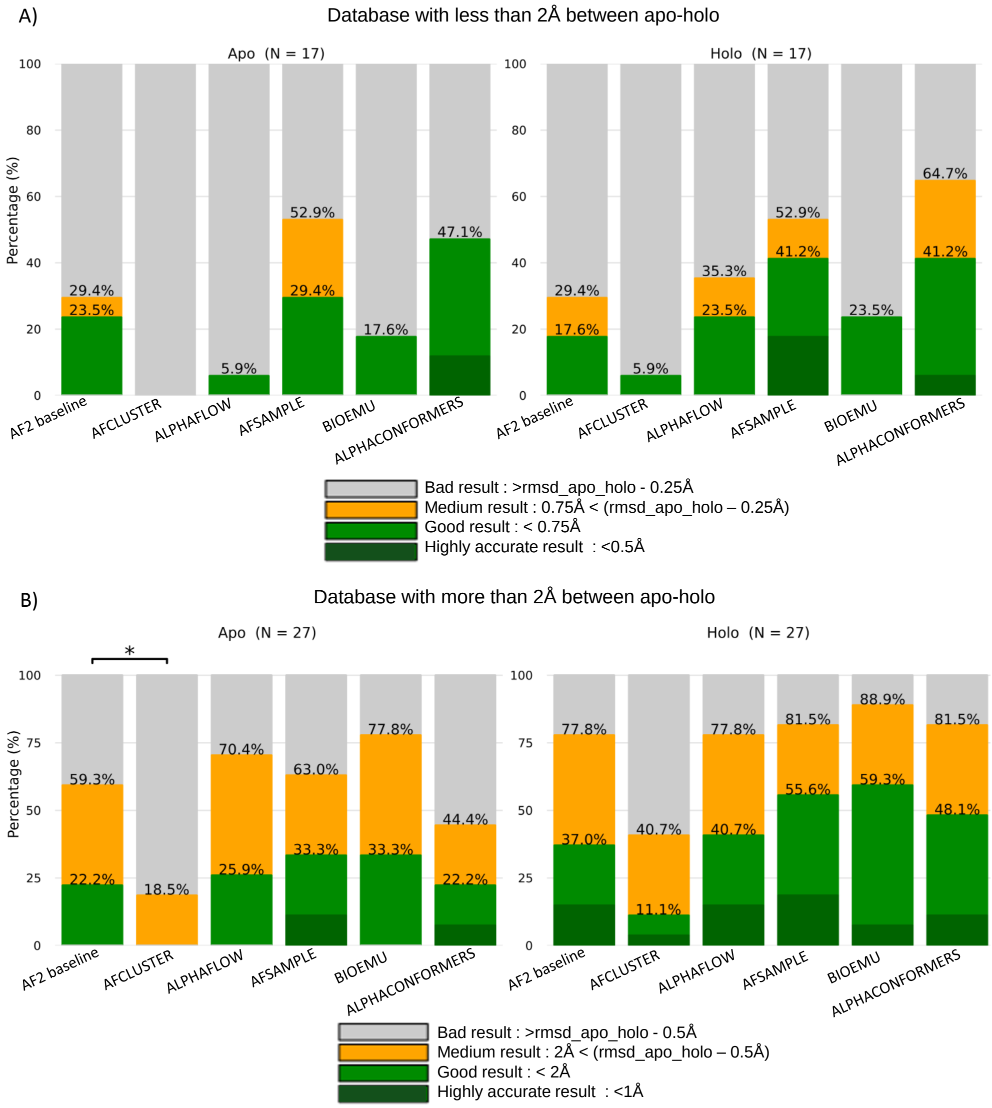


**Fig. S10: Benchmark comparison on the sequence homology AF2-exclusive subset,.** Restricted to proteins absent from AlphaFold2 training data with no structurally similar or homology sequence deposited prior to the AF2 training cutoff . **(A)** Recovery rates for the apo and holo states independently on the subtle-conformationa-change subset (apo-holo RMSD $< 2$ Å). AFsample3 and AlphaConformers outperform all other methods for apo and holo recovery. **(B)** Recovery rates for the apo and holo states independently on the large-conformational-change subset (apo-holo RMSD $\geq 2$ Å). For apo recovery, methods rank as follows: BioEmu and AFample3 (equal), AlphaFlow, AF2 baseline and AlphaConformers (equal), and AF-Cluster. For holo recovery: BioEmu, AFsample3 AlphaConformers, AlphaFlow, AF2 baseline and AF-Cluster.

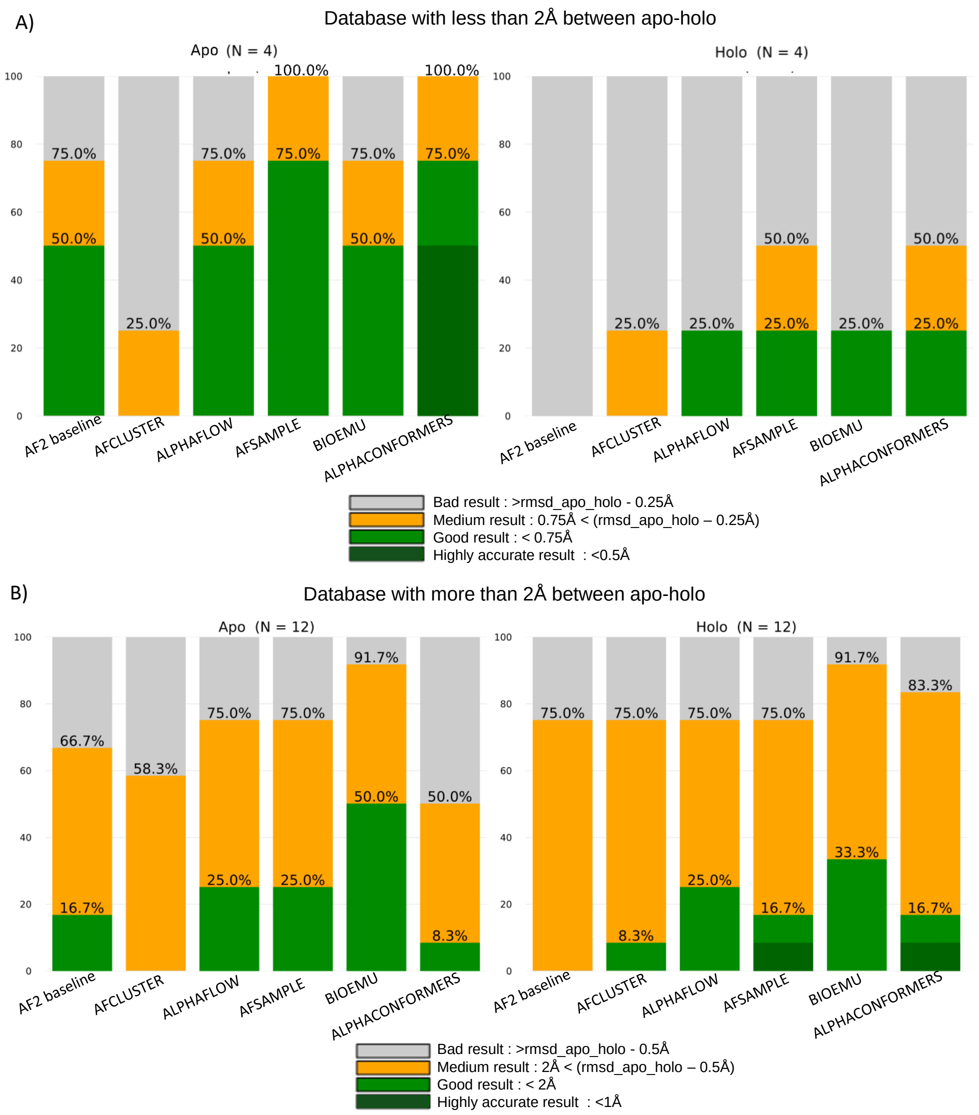


**Fig. S11: Benchmark comparison on the AF3-exclusive subset** Restricted to proteins absent from AlphaFold3 training data with no structurally similar entry deposited prior to the AF3 training cutoff.**(A)** Recovery rates for the apo and holo states independently on the subtle-conformational-change subset (apo-holo RMSD $< 2$ Å). AFsample3 and AlphaConformers outperform all other methods for apo recovery, while for the holo state all methods primarily contribute through conformational space exploration rather than exact recovery. **(B)** Recovery rates for the apo and holo states independently on the large-conformational-change subset (apo-holo RMSD $\geq 2$ Å). For apo recovery, methods rank as follows: BioEmu, AlphaFlow, AFsample3 (equal), AF2 baseline, AlphaConformers, and AF-Cluster. For holo recovery: BioEmu, AlphaFlow, AFsample3 and AlphaConformers (equal), AF-Cluster, and AF2 baseline.
